# NOX4 prevents the recruitment of PAX8 and NKX2.1 to chromatin in BRAF-mutated thyroid cancer cells

**DOI:** 10.1101/2025.04.14.647724

**Authors:** Mickaëlle Radom, Camille Buffet, Juliana Cazarin, Marylin Harinquet, Caroline Coelho de Faria, Floriane Brayé, Catline Nobre, Marine Aglave, Yasmina Mesloub, Thibault Dayris, Nathalie Droin, Karine Godefroy, Mohamed-Amine Bani, Abir Al Ghuzlan, Sophie Leboulleux, Livia Lamartina, Corinne Dupuy

## Abstract

Radioiodine (RAI) therapy, used for treating thyroid cancers, hinges on the expression of the Sodium Iodide Symporter (NIS). The majority of differentiated thyroid cancers (DTCs) are papillary, with a BRAF^V600E^ mutation. This mutation correlates with an absence of RAI uptake, due to low NIS expression and a low differentiation score. NADPH oxidase 4 (NOX4)-derived ROS contribute to NIS repression in BRAF^V600E^-mutated thyroid cancer cells. Depleting NOX4 enhances the reactivation of NIS. This reversibility implies an epigenetic mechanism’s contribution. Our findings indicate that NOX4 generates oxidative DNA damage in BRAF^V600E^-mutated thyroid cancer cells. DNA repair proteins such as OGG1 and MSH2/MSH6 proteins, in cooperation with DNMT1, turn these damages into transcription-blocking damages. This prevents the binding of PAX8 and NKX2.1 – two key transcription factors involved in thyroid differentiation – to the chromatin. Co-inhibition of the MAPK pathway, which regulates MSH2/MSH6 and DNMT1 expressions, and the TGF-β1 pathway, which regulates NOX4 expression, fortifies the recruitment of the two transcription factors to the chromatin. Collectively, our findings present a molecular basis for NOX4’s role in thyroid dedifferentiation.

## Introduction

Radioiodine therapy (RAI) relies on the capacity of thyroid cells to uptake and concentrate iodide. The sodium iodide symporter (NIS), situated in the basolateral membrane, mediates the active transport of iodide from the bloodstream into thyroid cells. The iodide is then processed by the unique iodide-metabolizing machinery, which is controlled by TSH, concentrates iodine into the cells, and has a significant impact on the efficacy of radioiodine [1]. Dedifferentiation correlates with a decrease or, in some cases, a complete loss of expression of thyroid-specific genes (such as *SLC5A5* (encoding NIS), *TPO*, *TG*, *TSHR*) and thyroid transcription factor genes (like *PAX8* and *NKX2.1* (TTF1). These last two genes play a critical role in regulating gene transcription activity in thyroid follicular cells [2].

A treatment approach for RAI-refractory patients is to re-enhance RAI uptake or re-differentiate tumors [1]. Indeed, restoring RAI uptake is the initial step of redifferentiation.

Most differentiated thyroid cancers (DTC) are papillary types (PTC), accounting for 80% of cases. These PTCs are often marked by the presence of the BRAF^V600E^ mutation in 45% to 60% of cases. This mutation is linked with aggressive tumor growth, low gene expression levels related to iodide metabolism, and low tumor differentiation scores [3] – all culminating in resistance to radioiodine (RAI) treatment. The precise mechanism behind this phenomenon remains obscure. BRAF is a potent activator of the MEK/MAP kinase pathway. Interestingly, in a designed mouse model, it was observed that inhibiting mutated BRAF or MEK resulted in reactivating NIS expression and thyroid tumor iodide uptake [4]. Nevertheless, the results varied in human studies [5]. Hence, deciphering the mechanisms behind the lowered expression of NIS and other thyroid function-related genes is crucial for establishing new molecular targets and developing novel treatment approaches.

Numerous studies have conclusively demonstrated that reactive oxygen species (ROS), via a variety of redox reactions, regulate cellular functions including gene expressions. In cells, NADPH oxidases (NOXs/DUOXs), which are membrane-bound complexes, are totally devoted to ROS production [6]. Three of these – DUOX2 (whose function is to provide H_2_O_2_ required for TPO) and DUOX1 and NOX4 (whose roles in this tissue are still unknown) – are expressed in thyroid cells. We previously demonstrated that ROS derived from NOX4 contributes to the repression of *SLC5A5* at the transcriptional level [7]. The expression level of NOX4 is significantly elevated in both human and murine BRAF-mutated thyroid tumors and is inversely correlated with thyroid differentiation. This suggests that genes involved in thyroid differentiation might be suppressed by a mechanism controlled by ROS derived from NOX4 [7].

The reversibility of the *SLC5A5* gene’s extinction by NOX4 suggests its contribution to an epigenetic mechanism, including gene methylation by DNA methyltransferases (DNMTs). DNA methylation plays a critical role in gene expression regulation, and recent findings have revealed a connection between oxidative damage repair by DNA mismatch repair (MSH2-MSH6) proteins and transcriptional inhibition by DNMT1 at DNA damage sites [8]. In this study, we demonstrate that NADPH oxidase NOX4, acting as an H_2_O_2_ generating system, leads to oxidative DNA damage in BRAF^V600E^-mutated thyroid cancer cells. These damages are transformed into transcription-blocking injuries by OGG1 and MMR proteins, which collaborate with DNMT1 and impede the binding of PAX8 and NKX2.1 (TTF1) to the chromatin, these are two key transcription factors involved in thyroid differentiation. Co-inhibition of the MAPK pathway and NOX4 enhances the recruitment of these two transcription factors to the chromatin. We also show that NOX4, OGG1, MSH2 and MSH6 are upregulated in Radioactive Iodine Refractory (RAIR) BRAF^V600E^-mutated thyroid cancer. Overall, our findings offer new insights into the role of NOX4-derived ROS in the thyroid dedifferentiation process.

## Results

### NOX4 is involved in oxidative DNA damage in BRAF-mutated thyroid cells

NOX4 is the only NOX with constitutive ROS-generating activity directly depending on its gene expression [9]. It is active in various intracellular compartments, including the nucleus. Compared to normal thyrocytes, NOX4 appears to be highly expressed in the nuclear fractions of three human cancer cell lines containing the BRAF^V600E^ mutation, with a higher expression in 8505C and the cell line established from Patient-Derived xenograft (PDX563) (Fig. 1a). The membrane protein, p22^phox^, serves as its functional partner and regulates both the stability and function of NOX4 [10]. RNAi-mediated depletion of either NOX4 or p22^phox^ in all BRAF^V600E^-mutated thyroid cell lines significantly decreases the nuclear H_2_O_2_ level, as evaluated by FACS using NucPE1, a nuclear-localized fluorescent H_2_O_2_ sensor (Supplementary Fig. 1a). Based on these results, we theorized that NOX4-dependent ROS production could be involved in oxidative DNA damage like the formation of 8-oxo-7,8-dihydroguanine (8-oxoG). The knockdown of either NOX4 or p22^phox^ significantly decreased the level of 8-oxoguanine as measured by cytometry in the three cell lines (Fig. 1b).

**Figure 1:**
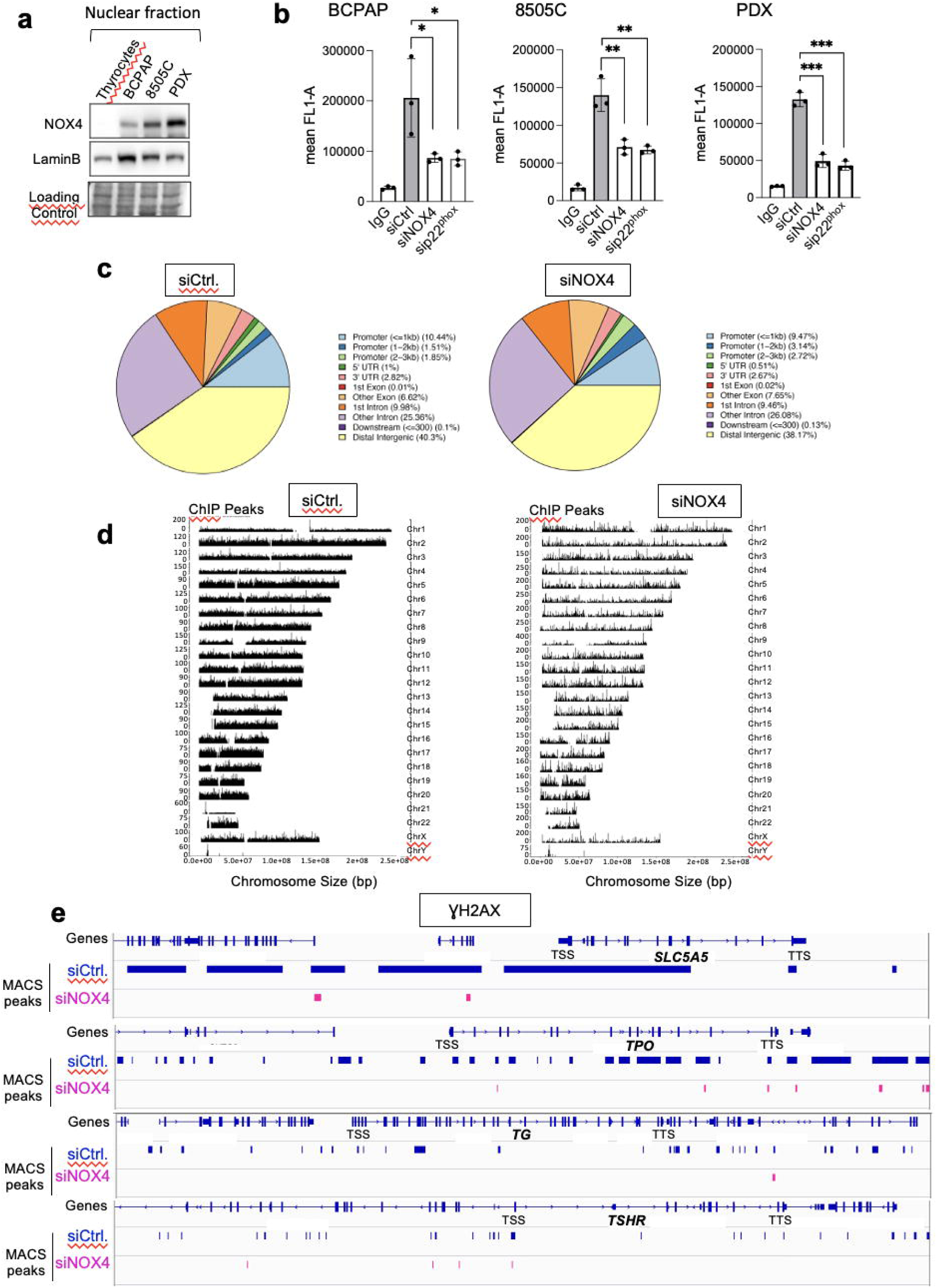
Knockdown of NOX4 or p22^phox^ reduces oxidative-DNA damage. **a**) Western-blot analysis of NOX4 protein expression in nuclear fractions of thyrocytes and BRAF-mutated thyroid cells. b) Quantification of 8-oxoG of BRAF-mutated thyroid cells by FACS analysis of immune-stained 8-oxoG cells. The cells were transduced with siRNA control or siRNA against NOX4 or p22^phox^ for 72 h. Graphs show the quantification of the fluorescence mean. Values are mean ± SE. *p < 0.05, **p < 0.01, and ***p < 0.001 (n = 3). c) Genomic distribution of γH2AX in BCPAP cells depleted or not for NOX4 (72h). Two replicates for each condition (siCtrl. and siNOX4) were pooled. d) Distribution of γH2AX on genome for BCPAP cells depleted or not for NOX4. The x axis corresponds to the position of the peaks along chromosomes; the y axis corresponds to the MACS2 score (-10*(log10(q_value)). e) γH2AX enrichment peaks on selected genes in conditions siRNA control and siRNA NOX4.

The phosphorylated form of histone H2AX ( γH2AX) at Ser^139^ is used as a marker of DNA damage. Next, we performed a Chromatin immuno-precipitation, followed by a high-throughput sequencing analysis of γH2AX distribution. Analysis of the genomic distribution of γH2AX peaks from biological replicates showed that they were localized within intergenic and intron regions (Fig. 1c). The mapping of γH2AX reads on the genome for BCPAP cells clearly showed that γH2AX enrichment was altered by RNAi-mediated NOX4 depletion (Fig. 1d). The sequence alignment analysis highlighted that γH2AX enrichment, occurring at genomic regions including the *SLC5A5* (NIS), *TPO*, *TG*, and *TSHR* genes, was impacted by this depletion (Fig 1e).

### NOX4-derived ROS mediates recruitment of MSH2, MSH6, and OGG1 to chromatin

8-oxodG is repaired either by the base excision repair (BER) pathway, a multistep process requiring several activated proteins like DNA glycosylase OGG1, or by the mismatch repair (MMR) system [11, 12]. The main mismatch-binding factor in human cells is MutS, which consists of a heterodimer of MSH2 and MSH6; it has been proven to play a role in repairing clustered oxidative DNA lesions [13]. To determine if MSH2, MSH6, and OGG1 are regulated by mutated BRAF, we transduced primary human thyrocytes with an expression vector that carries cDNA for BRAF^V600E^. Western-blot analysis revealed that overexpression of the activated oncogene caused an increase in the recruitment of MSH2 and MSH6 to the chromatin as well as in their protein levels (Fig. 2a and Supplementary Fig. 2a), but OGG1 was unaffected.

**Figure 2:**
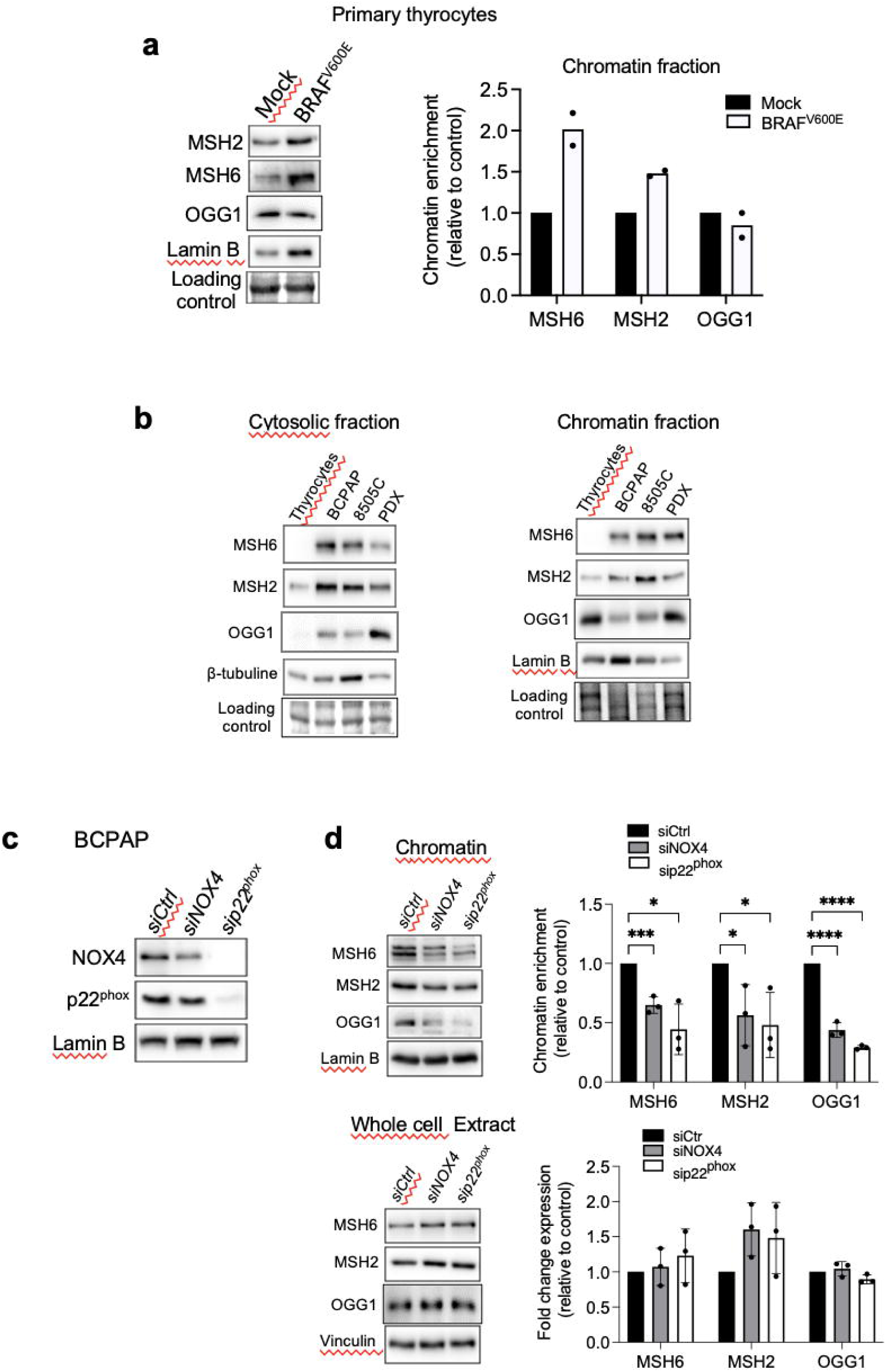
Knockdown of NOX4 or p22^phox^ reduces the recruitment of OGG1, MSH2, and MSH6 to chromatin. **a**) Human primary thyrocytes were transiently transfected with empty vector (Mock) or BRAF^V600E^ plasmids for 48 h and MSH2, MSH6, and OGG1 protein expression levels were analyzed by western-blot in chromatin fractions (n = 2). **b**) Western-blot analysis of MSH6, MSH2, and OGG1 protein expression levels in cytosolic and chromatin fractions of BRAF^V600E^-mutated thyroid cells. **c**) Western blot analysis of NOX4 and p22^phox^ protein expression levels in whole-cell extracts 72 h after knocking down of NOX4 or p22^phox^ by RNA interference in BCPAP cells. **d**) Western blot analysis of MSH6, MSH2, and OGG1 protein expression levels in chromatin fractions and whole-cell extracts 72 h after knockdown of NOX4 or p22^phox^ by RNA interference in BCPAP cells. Densitometry quantification of protein levels normalized to Lamin B or Vinculin levels and presented as chromatin enrichment or fold change compared with control cells. Values are mean ± SE. *p < 0.05, ***p < 0.001 and ****p<0.0001 (n = 3).

From TCGA data [14], we previously compared the mRNA expression levels of MSH2 and MSH6 in BRAF^WT^-PTCs (n = 170) and BRAF^MUT^-PTCs (n = 220). Both MSH2 and MSH6 showed a significant increase in PTCs containing a BRAF mutation [15]. Therefore, we compared the levels of MSH2, MSH6, and OGG1 proteins in both cytosolic and chromatin fractions of thyrocytes and BRAF-mutated thyroid cells (Fig. 2b). The levels of these DNA repair proteins were elevated in the cytosolic fractions of tumor cells compared to thyrocytes, with increased levels of MSH2 and MSH6 bound to chromatin. These findings suggest that BRAF^V600E^ regulates the expression and the recruitment of these proteins to chromatin in thyroid cancer cells..

Given the decrease in MSH2, MSH6, and OGG1 chromatin binding upon treating BRAF^V600E^ mutated thyroid cells with N acetyl cysteine (NAC), an antioxidant, or with Diphenylene Iodonium (DPI), an inhibitor of NADPH oxidases (Supplementary Fig. 2b and 2c), we studied their binding to chromatin after knocking down NOX4 and p22^phox^ (Fig. 2c and Supplementary Fig. 3a). Interestingly, NOX4 or p22^phox^ depletion significantly decreased the amount of MSH2, MSH6, and OGG1 bound to chromatin without altering their overall levels in the three cell lines (Fig. 2d, Supplementary Fig 3b and 3c, Supplementary Fig 4a and 4b). In addition, the affinity for chromatin of Myc-tagged OGG1 transfected into BCPAP and PDX cells decreased with NOX4 and p22^phox^ knockdown (Supplementary Fig 5a and 5b, Supplementary Fig 6a and 6b). These results collectively demonstrate the critical role of NOX4-derived ROS in increasing the recruitment of OGG1, MSH2, and MSH6 to chromatin in BRAF^V600E^-mutated thyroid cells.

### NOX4-derived ROS promotes the recruitment of DNMT1 protein to chromatin

The DNA methylases, DNMTs, are known as major mediators of DNA methylation. BRAF^V600E^ controls the expression of DNMT1 [16]. Consistent with this, DNMT1 was found to be elevated in the chromatin fractions of three human cancer cell lines harboring the BRAF^V600E^ mutation (Fig. 3a). Treating BRAF-mutated thyroid cells with NAC, DPI, or siRNA directed against NOX4 or p22^phox^ decreased the binding of DNMT1 to chromatin (Fig. 3b, Supplementary Fig 3a, Supplementary Fig 7a-7d). Conversely, the binding of DNMT3a and DNMT3b to the chromatin as well as their overall levels of expression were heightened in this condition (Supplementary Fig. 8a). This was reproduced when DNMT1 was depleted (Supplementary Fig 8b).

**Figure 3:**
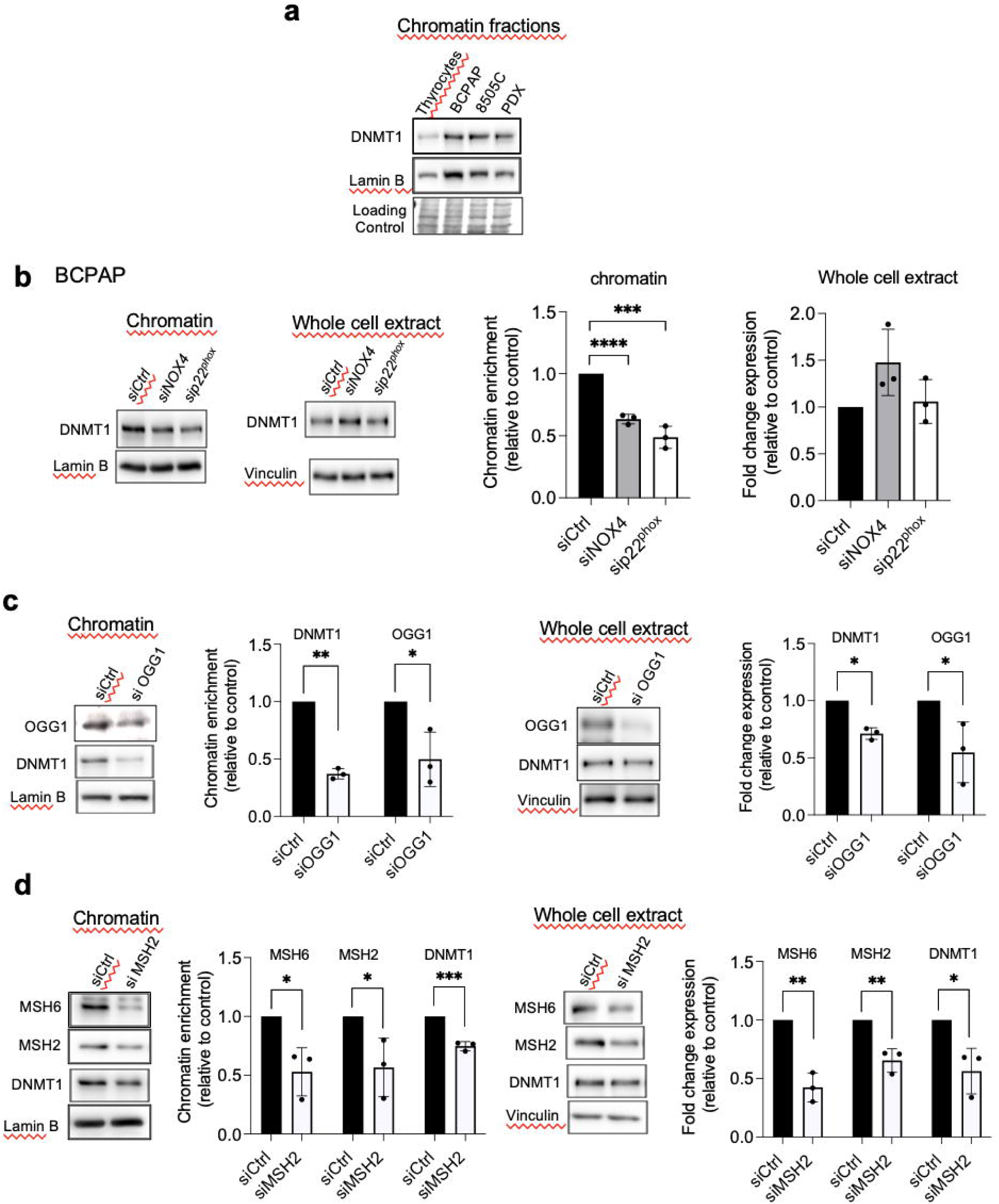
Knockdown of NOX4 or p22^phox^ as well as DNA repair proteins reduce the recruitment of DNMT1 to chromatin. **a**) Western-blot analysis of DNMT1 in chromatin fraction of thyrocytes and BRAF^V600E^-mutated thyroid cells. **b**) Western blot analysis of DNMT1 protein expression in chromatin fractions and whole-cell extracts 72 h after knocking down of NOX4 or p22^phox^ by RNA interference in BCPAP cells. **c**) Western blot analysis of DNMT1 protein expression in chromatin fractions and whole-cell extracts 72 h after knocking down of OGG1 by RNA interference in BCPAP cells. **d**) Western blot analysis of DNMT1 protein expression in chromatin fractions and whole-cell extracts 48 h after knocking down of MSH2 by RNA interference in BCPAP cells. Densitometry quantification of protein levels normalized to Lamin B or Vinculin levels and presented as chromatin enrichment or fold change compared with control cells. Values are mean ± SE. *p < 0.05, **p < 0.01, ***p < 0.001 and ****p<0.0001 (n = 3).

### OGG1 and MMR proteins are involved in the recruitment of DNMT1 to chromatin in BRAF-mutated thyroid cancer cells

The mismatch repair proteins MSH2 and OGG1 regulate the recruitment of DNMT1 to sites of oxidative DNA damage [8]. Thus, we further evaluated if these proteins regulate DNMT1 chromatin binding in BRAF-mutated thyroid cancer cells. Depletion of OGG1 or MSH2 decreases the binding of DNMT1 to the chromatin as well as its expression in whole-cell extracts (Fig. 3c and 3d). These findings suggest that these DNA repair proteins play a role in DNMT1 recruitment to areas of oxidative DNA damage produced by NOX4. As these results implied a role for NOX4 in DNA methylation, we analyzed the effects of NOX4 knockdown on genome-wide DNA methylation levels in BCPAP cells utilizing high-coverage nanopore sequencing. This technology, which allows for direct analysis of long DNA fragments without requiring PCR amplifications, enables simultaneous detection of nucleotide sequence and DNA base modifications, including 5-methylcytosine (5mC) on native DNA. Using this method, a maximum siRNA NOX4-induced methylation change of 50% was observed (Supplementary Fig 9a).

We then examined the genomic location of hypomethylated and hypermethylated regions following NOX4 deletion. Most were found to be enriched in intronic regions compared to promoters and exons (Supplementary Fig 9b). We selected 30 hypomethylated and hypermethylated regions with the highest differences for comparison between siRNA control and siRNA NOX4 (Supplementary Fig 9c). Among these genes, NOX4 depletion induced hypomethylation in exon 1 of *SMPD4* and exon 8 of *MTTL25* genes respectively, and it induced hypermethylation in the promoter region of PAX6 (Supplementary Fig 9d). This rise in hypomethylated regions suggests that NOX4 plays a role in recruiting DNMT1 to DNA methylation sites. Notably, no changes were seen in the proximal promoter regions of genes involved in thyroid differentiation, such as *SLC5A5*, *TSHR,* and *NKX2.1*, following NOX4 knockdown. These regions were found to be unmethylated in the BCPAP cell line (Supplementary Fig 9e). Additional analysis of the *SLC5A5* gene’s 5′ regions, which regulate NIS expression, such as the CpG-island2 known as “NIS distal enhancer” (-2152/-1887 from the ATG site) – previously found to be hypermethylated in thyroid tumors [17] – also showed no methylation (data not shown).

However, combining siRNA-mediated MSH2/MSH6 or OGG1 knockdown with Decitabine (DAC), a DNMT inhibitor, resulted in a significant increase in the reactivation of the *SLC5A5* gene. This indicates that MMR and OGG1 proteins cooperate with DNMT1 to maintain the silencing of this gene (Fig. 4a). We were able to reproduce these results by combining siRNA-mediated NOX4 or p22^phox^ depletion with DAC (Fig. 4b). These results, taken together, suggest that NOX4 mediates the recruitment of MMR, OGG1, and DNMT1 proteins on damaged chromatin and these may work synergistically to repress genes such as *SLC5A5*.

**Figure 4:**
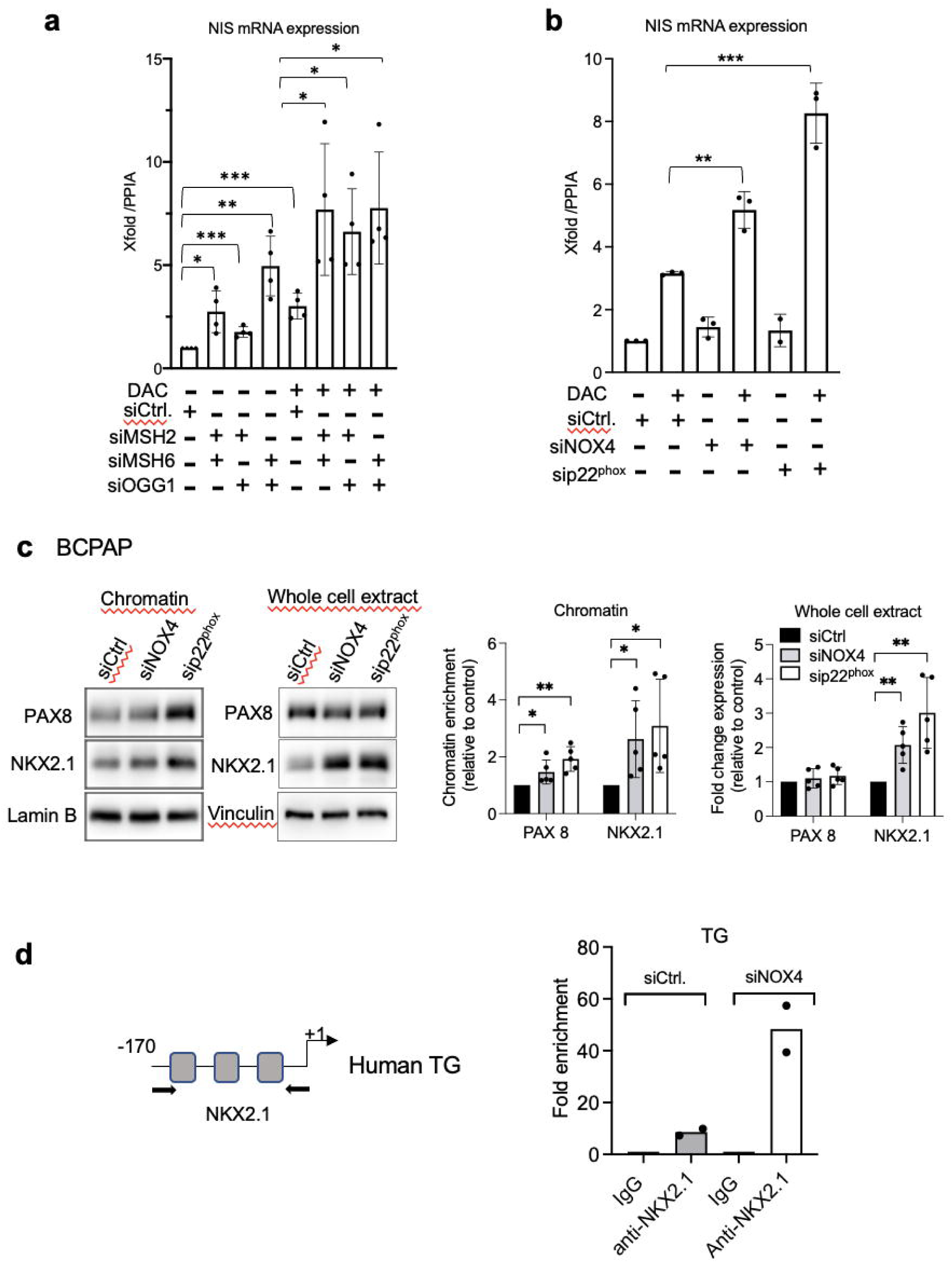
DNMT inhibition and knockdown of MSH2, MSH6, and OGG1 synergize to increase NIS mRNA expression. **a**) qRT-PCR analysis of NIS mRNA levels in BCPAP cells transfected with siRNA control or/and siRNA MSH2 or/and siRNA MSH6 or/and siRNA OGG1 and 24 h later treated for an additional 48 h in the presence or the absence of 1 µM DAC. **b**) qRT-PCR analysis of NIS mRNA levels in BCPAP cells transfected with siRNA control or siRNA NOX4 or siRNA p22^phox^ and 24 h later, treated for an additional 48 h in the presence or the absence of 1 µM DAC. Values are mean ± SE. *p < 0.05, **p < 0.01, and ***p < 0.001. **c**) Western blot analysis of PAX8 and NKX2.1 protein expressions in chromatin fraction and whole-cell extract 72 h after knocking down of NOX4 or p22^phox^ by RNA interference in BCPAP cells. Densitometry quantification of protein levels normalized to Lamin B or Vinculin levels and presented as chromatin enrichment or fold change compared with control cells. Values are mean ± SE. *p < 0.05 and **p < 0.01 (n = 3). **d**) ChIP-qPCR assays performed with BCPAP cells transfected with siRNA control and with BCPAP cells transfected with siRNA NOX4 immunoprecipitated with control IgG or anti-NKX2.1 antibody and analyzed by qPCR at TG promoter (two independent replicates).

### NOX4-derived ROS prevents the recruitment of transcription factors NKX2.1 and PAX8

The promoters of the primary thyroid differentiation markers are regulated by the specific combination of the transcription factors NKX2.1 and PAX8 [2]. A transient induction in human embryonic stem cells enables efficient generation of thyroid follicular cells capable of thyroid hormone production [18]. NAC or DPI treatment of BRAF-mutated thyroid cells increased the binding of PAX8 and NKX2.1 to chromatin, as well as their total level of expression, indicating that the recruitment and expression of these proteins are redox-sensitive (Supplementary Fig. 10a and 10b). We then analyzed the inhibitory effect of NOX4 on these proteins’ binding to the chromatin. Importantly, the knockdown of NOX4 or p22^phox^ significantly increased the affinity of both transcription factors for chromatin in the analyzed BRAF-mutated thyroid cell lines BCPAP and PDX (Fig. 4c, Supplementary Fig 10c and Supplementary Fig 11a). Since PAX8 was undetectable in 8505C cells, only the effect on NKX2.1 was analyzed, and the same result was obtained (Supplementary Fig 11b). Three sites on the TG proximal promoter are recognized by NKX2.1 [19]. Using a ChIP qPCR assay, we observed that the knockdown of NOX4 increased NKX2.1 binding to the promoter (Fig. 4d).

Also, the knockdown of OGG1 or both MSH2 and MSH6 facilitated the recruitment of the two transcription factors to the chromatin in the BRAF-mutated cell lines (Fig 5a, Supplementary Fig 12a-12c). Depending on the cell line, their total level of expression also appeared to be modulated (Supplementary Fig 13a and 13b, Supplementary Fig 14a-14c). These results suggest that NOX4-derived ROS prevent PAX8 and NKX2.1 from binding to the chromatin through the recruitment of DNA repair proteins to the chromatin, induced by oxidative damage.

**Figure 5:**
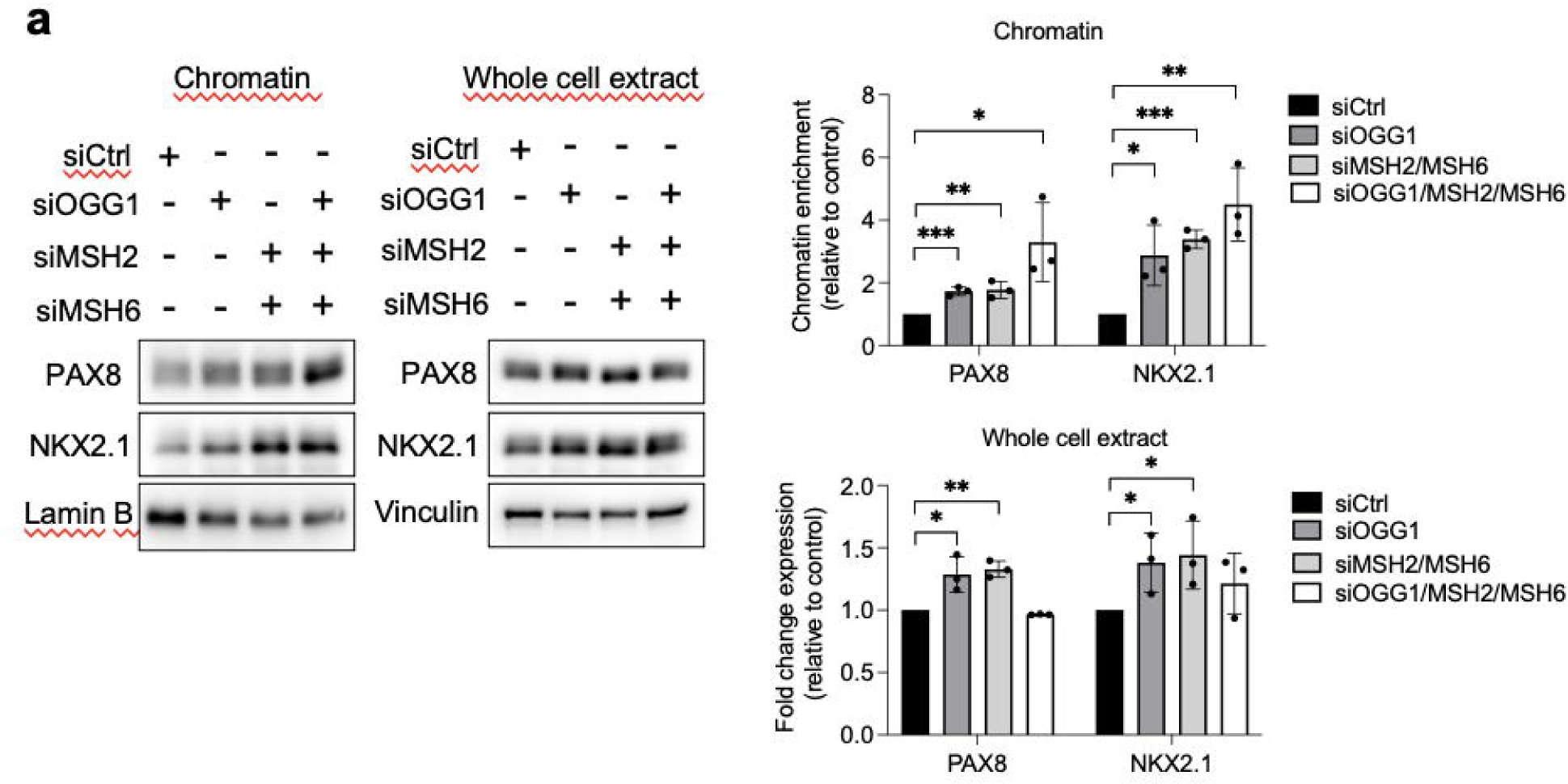
OGG1, MSH2, and MSH6 inhibit recruitment of PAX8 and NKX2.1 to chromatin. **a**) Western blot analysis of PAX8 and NKX2.1 protein expressions in chromatin fraction and whole-cell extract 72 h after knocking down of OGG1 (siRNA OGG1#1), MSH2, or/and MSH6 by RNA interference in BCPAP cells. Densitometry quantification of protein levels normalized to Lamin B or Vinculin levels and presented as chromatin enrichment and fold change compared with control cells. Values are mean ± SE. *p < 0.05, **p < 0.01, and ***p < 0.001 (n = 3).

To further investigate the influence of NADPH oxidase on genome-wide chromatin accessibility, we performed an Assay for Transposase-Accessible Chromatin using sequencing (ATAC-seq) on BCPAP, 8505C, and PDX563 tumor thyroid cells, with or without p22^phox^ depletion (Supplementary Fig 15a-15e and Supplementary Fig 16a-16f). We detected a relatively distinct footprint of TF occupancy near the aggregated PAX8 and NKX2.1 full sites in accessible chromatin regions in all control and p22^phox^-depleted cells. Moreover, the compaction level of PAX8 and NKX2.1 genes was not altered by p22^phox^ depletion. Intergenic and intron regions were the most enriched genomic features of the accessible regions in the BRAF-mutated thyroid cell lines. Most of the peaks were 0-1 kb and 10–100 kb away from TSS, a distribution not affected by p22^phox^ depletion. Lastly, the abrogation of p22^phox^ expression did not modulate accessibility to the set of thyroid differentiation marker genes.

### MSH2 and MSH6 expressions are inhibited by the combination of dabrafenib/trametinib

Clinical evidence has shown a link between MAPK pathway inhibition and increased molecular differentiation in patients, supporting that MAPK is a critical regulator of thyroid tumor differentiation [1]. The combination of dabrafenib/trametinib inhibits both mutated BRAF and MEK in all cell lines, demonstrated by the inhibition of phospho-MEK and phospho-ERK (Supplementary Fig 17a). A 72-h treatment of BRAF-mutated thyroid cell lines with this combination led to a downregulation of MMR proteins, indicating that the MAPK pathway controls the expression of these proteins in tumor cells (Fig. 6a and Supplementary Fig. 17b and 17c). Interestingly, while this treatment altered the DNMT1 protein level, it had the opposite effect on the OGG1 protein level, which was upregulated in this condition (Fig. 6b and Supplementary Fig. 18a).

**Figure 6:**
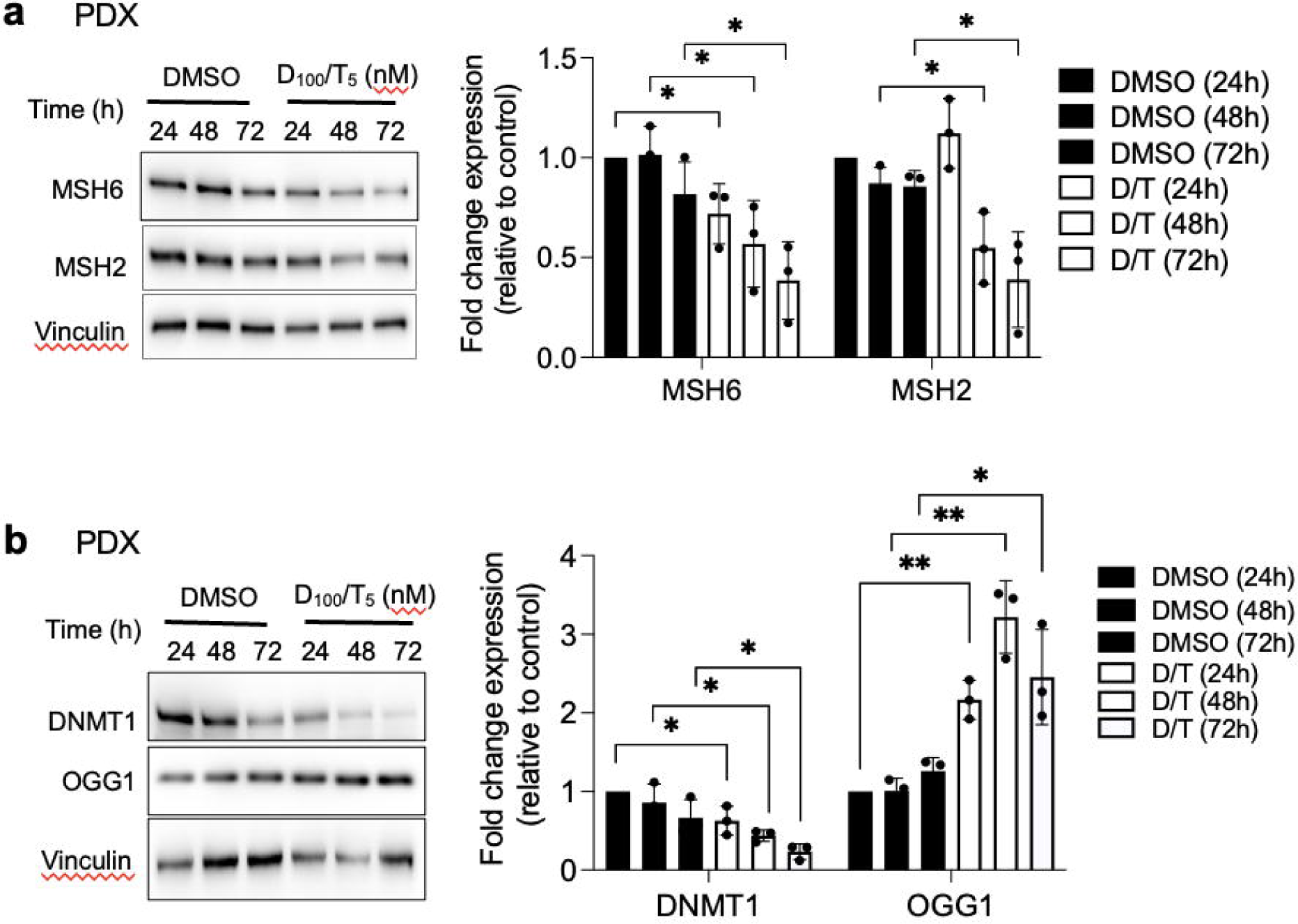
BRAF^V600E^ regulates MSH2 and MSH6 expressions. **a, b**) Immunoblot detection of MSH2, MSH6 DNMT1, and OGG1 in PDX cells after treatment by the combination of dabrafenib (100 nM) plus trametinib (5 nM). Densitometric quantification of protein levels normalized to vinculin levels and presented as fold change compared with vehicle-treated cells. Student t-test is realized by comparing combination versus DMSO for each corresponding time of kinetic. Values are mean ± SE. *p < 0.05 and **p < 0.01 (n = 3).

### NOX4 cooperates with the MAPK pathway in the thyroid dedifferentiation process

The combined knockdown of NOX4 or p22^phox^, in conjunction with dabrafenib/trametinib treatment, led to increased recruitment of NKX2.1 and PAX8 to the chromatin (Fig. 7a and Supplementary Fig. 18b-18c). This was associated with an increase in the expression of NIS and TSHR at the mRNA levels, which indicates that the NADPH oxidase NOX4 cooperates with the MAPK pathway in the thyroid dedifferentiation process (Fig. 7b and 7c).

**Figure 7:**
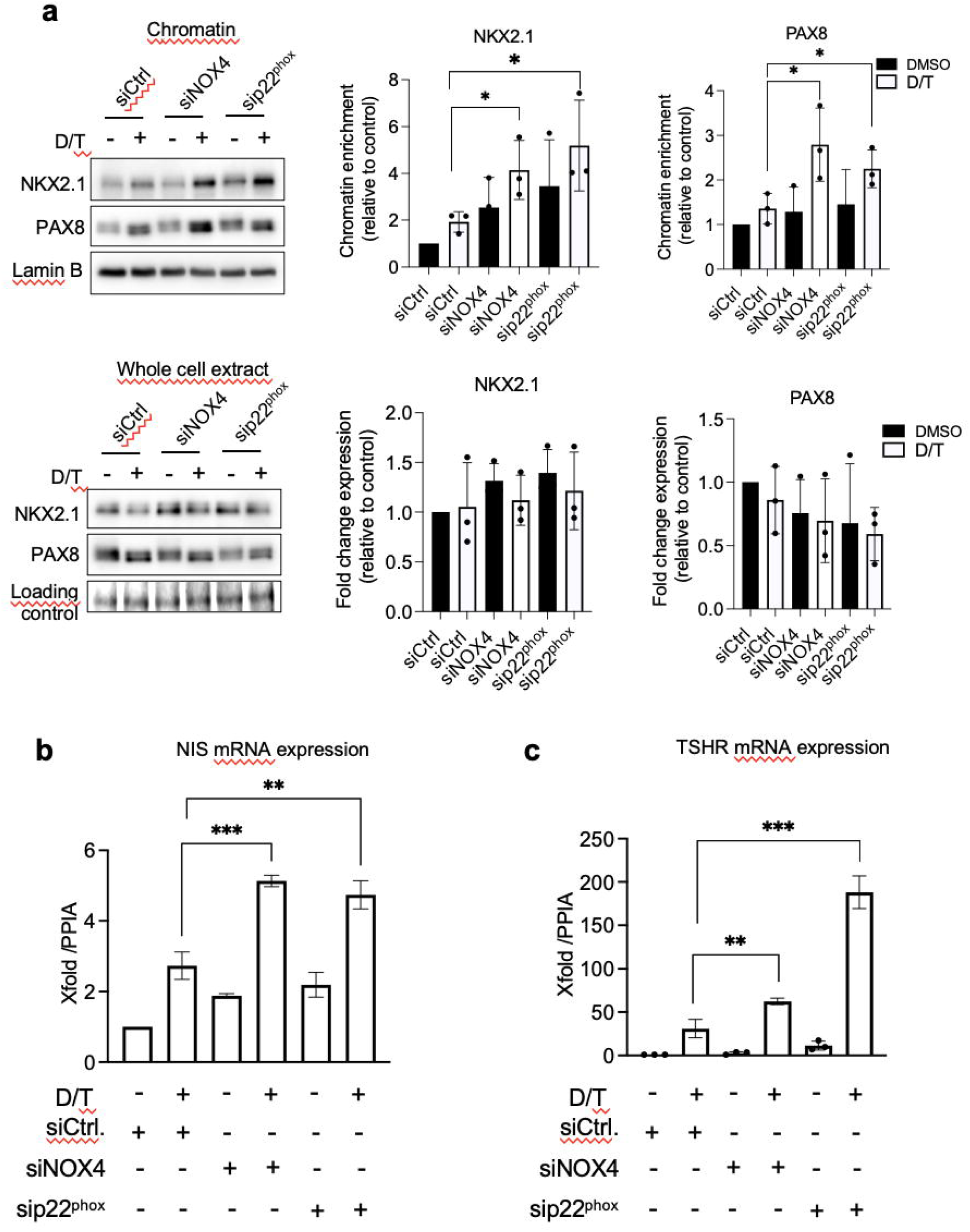
MAPK pathway inhibition and knockdown of NOX4 and p22^phox^ synergize to promote PAX8 and NKX2.1 recruitment to chromatin. **a**) Western-blot analysis of PAX8 and NKX2.1 protein expressions in chromatin fractions and whole-cell extracts of BCPAP cells transduced with siRNA control or siRNA NOX4 or siRNA p22^phox^ and treated with dabrafenib (100 nM) plus trametinib (25 nM) combination for 48 h. Densitometric quantification of protein levels normalized to Lamin B or loading control and presented as chromatin enrichment and fold change compared with siRNA control-transduced cells. **b**) qRT-PCR analysis of NIS mRNA levels in BCPAP cells transfected with siRNA control or siRNA NOX4 or siRNA p22^phox^ and 24 h later, treated in the presence or the absence of dabrafenib plus trametinib combination for additional 48 h. **c**) qRT-PCR analysis of TSHR mRNA levels in BCPAP cells transfected with siRNA control or siRNA NOX4 or siRNA p22^phox^ and 24 h later, treated in the presence or the absence of dabrafenib plus trametinib combination for additional 48 h. Values are mean ± SE. *p < 0.05, **p < 0.01, and ***p < 0.001 (n = 3).

We have previously shown that the TGF-β1 signaling pathway upregulates NOX4 in BRAF-mutated thyroid cells at the transcriptional level in a Smad3-dependent manner [7]. Luckett et al. used a genetically engineered mouse model of BRAF mutant thyroid cancer to show that suppressing both the MAPK and pSMAD pathways led to enhanced radioiodine uptake in mouse cancer cells [20]. Considering these results, we evaluated the combined effect of Dabrafenib/Trametinib and Vactosertib (EW7197), an inhibitor of the TGF-β1 pathway, on human BRAF mutated thyroid cancer cells. EW7197 increased the recruitment of PAX8 and NKX2.1 to the chromatin, and this effect was potentiated when the inhibitor was combined with MAPK kinase pathway inhibitors. This led to an increase in both TSHR and NIS mRNA expressions (Fig. 8a-8c, Supplementary Fig. 19a and 19b).

**Figure 8:**
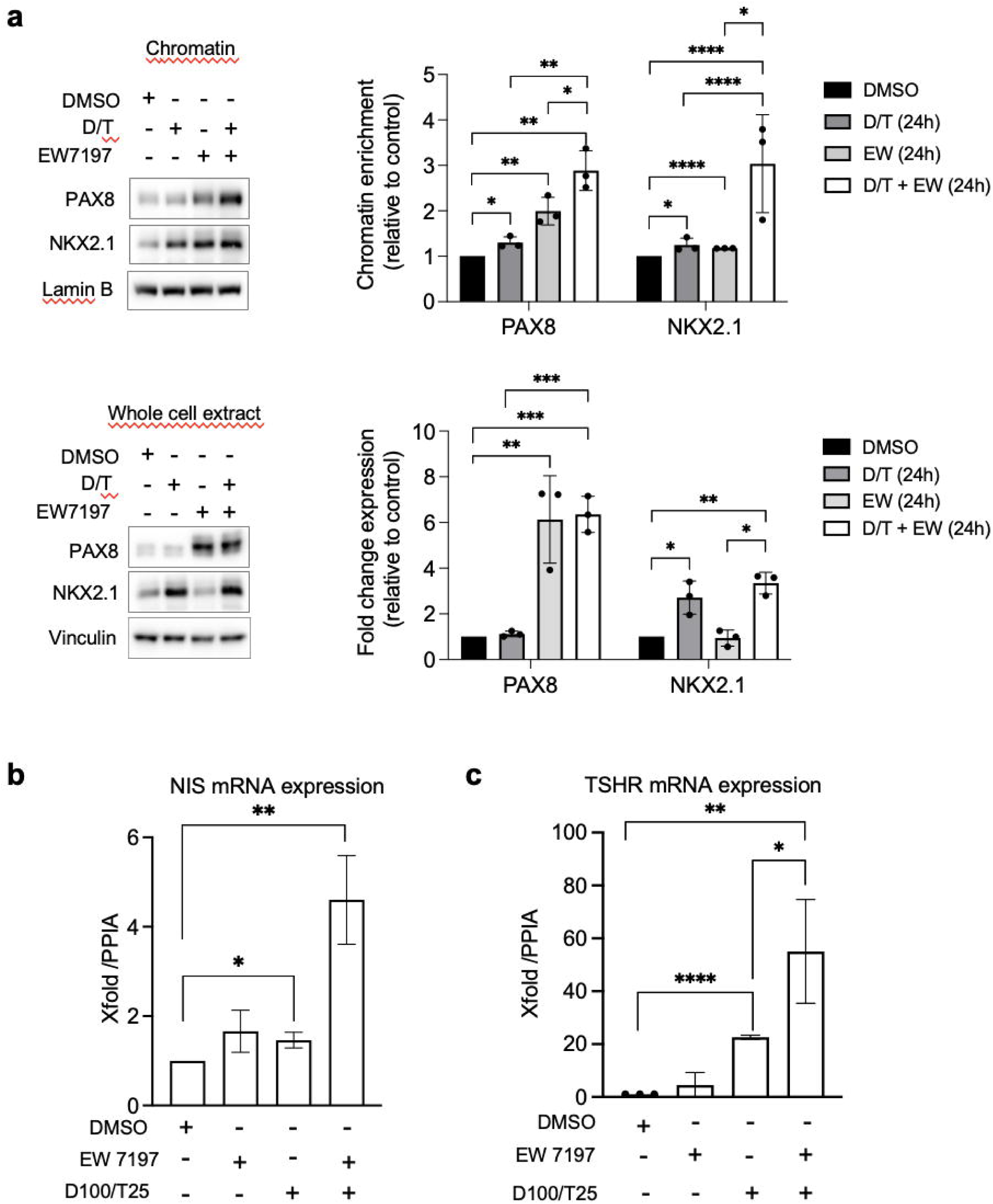
Co-inhibition of SMAD and MAPK signaling promotes PAX8 and NKX2.1 recruitment to chromatin. **a**) Western-blot analysis of PAX8 and NKX2.1 protein expressions in chromatin fractions and whole-cell extracts of BCPAP cells treated or not with dabrafenib (100 nM) plus trametinib (25 nM) combination in the presence or the absence of TGF-beta inhibitor (EW7197, 1 µM) for 48 h. Densitometric quantification of protein levels normalized to loading control and presented as chromatin enrichment or fold change compared with vehicle-treated cells. **b**) qRT-PCR analysis of NIS and TSHR mRNA levels in BCPAP cells treated or not with dabrafenib plus trametinib combination in the presence or the absence of TGF-beta inhibitor (EW7197) for 48 h. Densitometry quantification of protein levels normalized to Lamin B or Vinculin levels and presented as fold change compared with control cells. Values are mean ± SE. *p < 0.05, **p < 0.01, ***p < 0.001 and ****p<0.0001 (n = 3).

### NOX4, OGG1, MSH2, MSH6 and phospho-Smad3 are upregulated in Radioactive Iodine Refractory (RAIR) BRAF^V600E^-mutated thyroid cancer

A phase II redifferentiation trial with dabrafenib/trametinib and ^131^I in metastatic radioactive iodine refractory BRAF^V600E^-mutated differentiated thyroid cancer was previously conducted to explore therapeutic efficacy of drug combination [21]. The patients (n=21) underwent 6-weeks of treatment with dabrafenib and trametinib followed by a therapeutic activity of RAI (5.5 GBq of ^131^I) administered following two rhTSH injections. Some patients displayed primary resistance to this redifferentiation strategy and one patient achieved a complete response. From 15 of primary tumor tissue of these patients we undertook to analyze NOX4, DNA repair proteins (OGG1, MSH2 and MSH6) and phospho-Smad3 expressions by immunohistochemistry (Supplementary Table 1). Compared to normal tissue we identified an increased expression of NOX4, OGG1, MMR proteins and phosphor-Smad3 in RAIR BRAF^V600E^ mutated tumors (Fig. 9a). In contrast the two transcription factors PAX8 and NKX2.1 showed the same level of expression. We identified a significant positive correlation between MSH6 and OGG1 expression (p=0.0415, Spearman r=0.5361) and between phospho-Smad3 and NOX4 (p=0.0219, Spearman r=0.5926)

**Figure 9:**
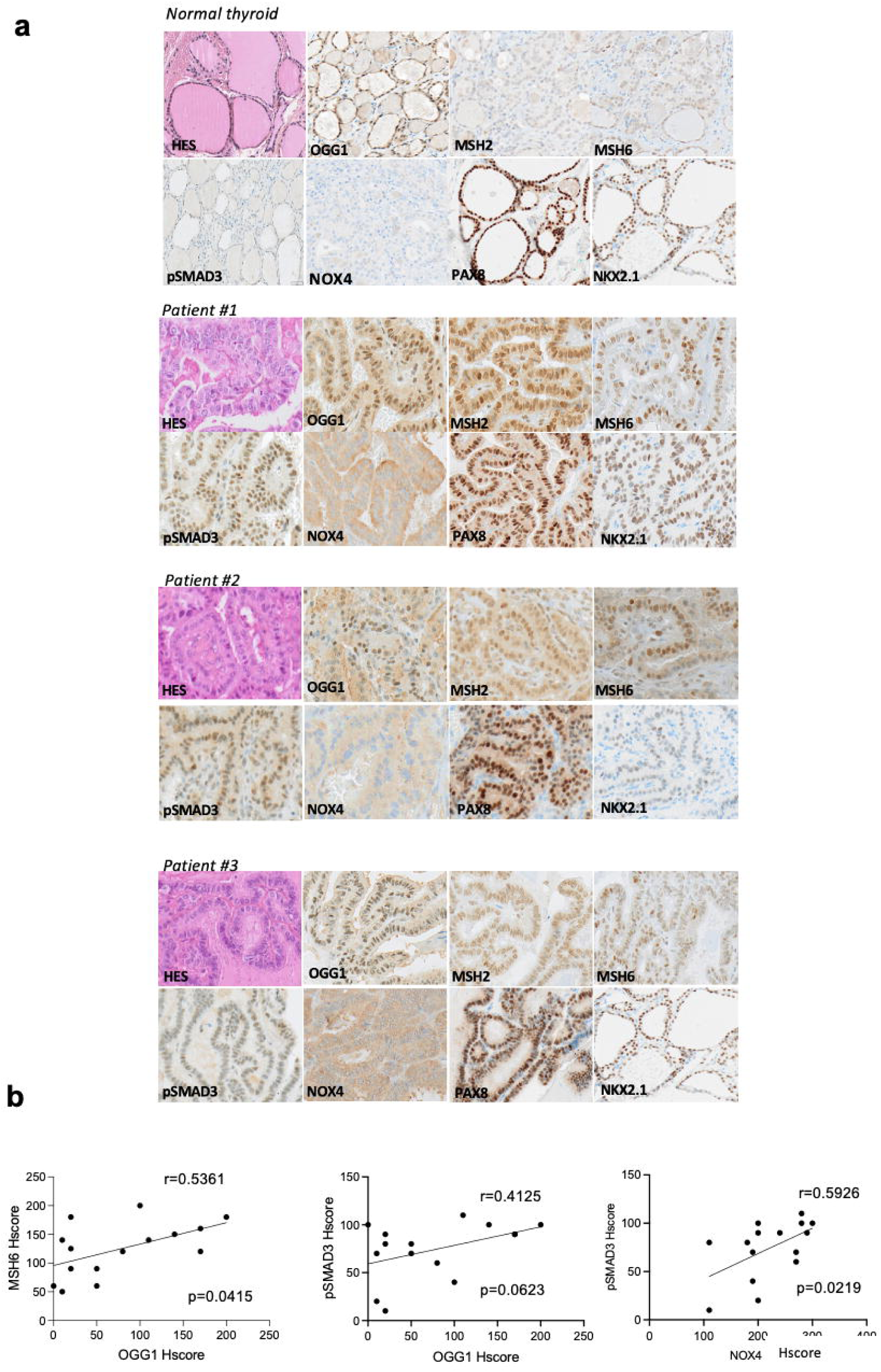
NOX4, oxidative DNA repair proteins and phospho-Smad3 are upregulated in radioactive iodine refractory BRAF^V600E^-mutated thyroid cancer. **a**) Immunoexpression of the different markers in normal thyroid tissue and in BRAF-mutated thyroid tumors. **b**) Correlation between immunohistochemical markers.

## Discussion

The findings reported here provide new insights into our understanding of the epigenetic role of 8-oxoG in BRAF^V600E^-mutated thyroid cancers, emphasizing the contribution of oxidative DNA damage in redox-mediated control of gene expression in thyroid dedifferentiation processes by altering the recruitment of two key transcription factors, PAX8 and NKX2.1.

It has been previously demonstrated that oxidative damage, induced by treating cancer cells with 1 mM hydrogen peroxide (H_2_O_2_), results in the localization of epigenetic silencing complexes to GC-rich areas of the genome, leading to decreased gene expression [22]. This subsequently raised questions regarding the critical role of endogenous H_2_O_2_-generating enzymatic systems in epigenetic mechanisms associated with oxidative DNA damage, particularly considering that H_2_O_2_ is not uniformly distributed within cells and its production induces DNA damage in a dose-dependent manner [23]. NADPH oxidases are one of the primary sources of ROS production in human cells, where they regulate physiological redox-dependent processes. Notably, NOX4, particularly active in nuclei, is associated with disease progression [24].

Given our previous discovery that NOX4 expression increases in BRAF^V600E^-mutated thyroid tumors, we aimed to study its role in an epigenetic mechanism contributing to the repression of genes related to thyroid differentiation. In the current study, using three BRAF^V600E^-mutated cell lines, including one derived from a patient-derived xenograft, we determined that BRAF-mutated thyroid cells express NOX4-derived H_2_O_2_ in their nuclear environment and that it is responsible for the formation of 8-oxoG DNA damage.

8-oxoG, in conjunction with OGG1, can influence transcriptional regulation (either activation or repression) by manipulating transcription factor homing or the recruitment of chromatin remodelers [25]. A significantly higher expression of the OGG1 gene and its corresponding protein has recently been noted in lesions of papillary thyroid cancer (PTC) compared to benign lesions [26]. OGG1 is drawn to open chromatin regions where 8-oxoG effectively recruits BER enzymes [27]. However, OGG1 cannot excise 8-oxoG within clustered lesions. Alternate DNA repair systems such as the non-canonical Mismatch Repair (MMR) system, including proteins MSH2 and MSH6, manage 8-oxoG in these situations. This is particularly pertinent in the repair of clustered oxidative damage found in GC-rich regions of the genome, including promoter CpG islands [13]. Higher levels of MSH2 have been identified in malignant thyroid tumors in comparison to benign tumors [28]. In the present work we show that both OGG1 and MMR proteins are up-regulated in iodine-refractory BRAF^V600E^-mutant metastatic differentiated thyroid cancer, indicating a functional activation of these genes. Thyroid tumors exhibit a low overall density of somatic mutations compared to other tumors, suggesting efficient support from both the BER and MMR systems. Moreover, the loss of MSH2 and MSH6 was associated with a hypermutated phenotype found in anaplastic thyroid carcinoma, a highly aggressive and undifferentiated malignant tumor [29]. However, the involvement of MMR proteins in the development and progression of thyroid cancer still needs to be demonstrated. Knockdown of NOX4 or its functional partner p22^phox^, as well as antioxidant treatment, inhibited the recruitment of both OGG1 and MMR proteins to the chromatin in BRAF^V600E^-mutated thyroid cells. Also, the depletion of these DNA repair proteins independently reactivated the expression of the *SLC5A5* gene, which encodes NIS. These findings suggest that NOX4-derived 8-oxoG can be converted into transcription-blocking damage by OGG1 and MMR proteins in BRAF^V600E^-mutated thyroid cancer cells.

Evidence is showing that OGG1 may play multiple role [30]. Thus, it also promotes the recruitment of transcription factors, such as NF-κB, and the assembly of transcriptional machinery to assure a prompt launch of the immediately responsive transcriptome. Whether NOX4-derived 8-oxoG promotes enrichment of NF-κB on promoters of proinflammatory genes needs to be further studied, especially since NOX4 was shown to regulate the expression of cytokines in several cancers.

The pattern of DNA methylation is altered in thyroid cancer. Papillary thyroid cancer (PTC) manifests one of the lowest frequencies of DNA methylation [31]. The BRAF^V600E^ mutation is more commonly associated with the classical subtype of PTC and correlates with an increase in DNA hypomethylation, a trend observed in other malignancies largely dependent on stable maintenance by DNMT1 [32]. BRAF^V600E^ regulates DNMT1, whose expression is elevated in PTCs that express this oncogene [16]. Our findings indicate that the hindrance of DNMT1 binding to the chromatin due to NOX4 knockdown coincides with alterations in the cell’s DNA methylation profile, causing some regions to become hypomethylated and others hypermethylated.

A recent study spotlighted the cooperative activity between DNMT1 and DNMT3b in maintaining and controlling DNA methylation using a DNMT1 degradation system [33]. Upregulation of DNMT3b was observed under DNMT1 degradation. Interestingly, our observations reveal that the knockdown of NOX4, as well as OGG1 or MSH2, which all affect DNMT1’s recruitment to the chromatin, led to an increase in both the expression and recruitment of DNMT3a/b to chromatin. This suggests that these DNMTs may perform a compensatory role at selected loci when DNMT1 levels diminish. As NOX4 is active at the nuclear membrane, this implies that the changes in DNA methylation observed in its absence might be preferentially located at the nuclear periphery. Abnormal methylation of promoter regions of thyroid marker genes, such as *TSHR* and *SLC5A5*, has been reported in PTC tumors, although the results for *SLC5A5* are controversial [31]. These regions were found to be unmethylated in BCPAP cells, but combining siRNA-mediated MSH2/MSH6 or OGG1 depletion or either siRNA-mediated NOX4 or p22^phox^ depletion with Decitabine, a DNMT inhibitor, resulted in a drastic reactivation increase of the *SLC5A5* gene. DNMT1 is necessary to maintain CpG methylation, but recent data suggest it can also modulate gene expression independent of its catalytic activity and participate in multiple processes, including the cell cycle and DNA damage repair [34]. Overall, our findings suggest NOX4-mediated recruitment of OGG1 and MMR proteins on damaged chromatin, which may synergistically cooperate with DNMT1 to maintain the silencing of genes.

NKX2.1 (or TTF1) and PAX8 are integral to thyroid organogenesis and the continued differentiation of thyrocytes [2]. Previous research revealed a redox regulation of Pax8 and NKX2.1 is implicated in their heightened DNA-binding activities prompted by thyrotropin in rat thyroid FRTL-5 cells [35]. Our findings indicate that ROS, sourced from NOX4, and by facilitating the engagement of OGG1 and MMR proteins with damaged chromatin, impacts the engagement of these two transcription factors with chromatin in BRAF-mutated thyroid cancer cells. Thus, this contributes to the process of thyroid dedifferentiation. Interestingly, PAX8 and NKX2.1 are expressed in the radioactive iodine refractory BRAF^V600E^-mutated thyroid tumours analyzed, highlighting that a decrease in their binding to chromatin and not in their level of expression may be one of the causes of the refractoriness of these tumors.

Loss of differentiation features correlates with the degree of MAPK activation, which is higher in tumors with the BRAF^V600E^ mutation. The demonstration that MAPK pathway inhibition restores the expression of genes that mediate iodide uptake has renewed interest in redifferentiation strategies [1, 4]. The combination of Dabrafenib/Trametinib synergistically increased iodide uptake in human BRAF-mutated thyroid cancer cell lines [36]. Here we demonstrate that BRAF^V600E^ controls the expression of MSH2 and MSH6 through an ERK-dependent pathway, providing an additional element to the understanding of the efficacy of inhibitors of the MAPK-signaling pathway in clinic. Along with other DNA repair genes, these MMR proteins increase in the presence of the E2F transcription factors, which the ERK cascade then controls through the regulation of Rb protein phosphorylation, one of their inhibitors [37]. Contrarily, MAPK pathway inhibition, surprisingly, leads to increased OGG1 expression. Many tumors that initially respond to therapy eventually develop resistance. Changes in DNA repair gene expressions are one possible cause of this resistance. This suggests that OGG1, by promoting DNA repair in tumor cells, contributes to therapy resistance in some ways. Further investigation of this is required.

The inhibition of MAPK transcriptional output did not result in sufficient redifferentiation in all treated patients. This suggests that more potent pathway inhibition may be required, or that biological factors beyond MAPK inhibition could be critical. Our data demonstrates that the knockdown of NOX4 or p22^phox^ enhances the effect of the drug combination on the recruitment of PAX8 and NKX2.1 to the chromatin. This is correlated with a higher reactivation of the *SLC5A5* and *TSHR* genes. Previously, it was established that the activation of the RAF/MEK/ERK pathway only accounts for a portion of NKX2.1 (TTF1) inactivation [38]. In this study, we identify NOX4 as an additional effector working in tandem with the RAF/MEK/ERK cascade to fully repress the functions of NKX2.1 (TTF1) and PAX8.

In general, antioxidants, such as NAC, have failed to demonstrate efficacy in the clinic. NAC is a general scavenger of ROS and, as such, can cause adverse cellular effects and unwanted toxicity because of the relevance of ROS to normal physiological functions. Thereby, effective treatment has to require maintaining a delicate balance between suppressing deleterious effects of ROS and interfering with cellular physiology, such as thyroid hormone biosynthesis, which is dependent of DUOX2-derived H_2_O_2_. Therefore, targeting the primary enzymatic source of ROS, or the signalling pathway that controls it, may have better therapeutic efficacy and reduce undesired side effects.

A TGF-β1 transcriptional output signature was identified in advanced RAI-refractory human BRAF-mutant thyroid cancers [20]. TGF-β1 was first demonstrated to play a significant role as a local thyroid modulator by inhibiting both growth and differentiation in various species [39]. The expression of BRAF^V600E^ triggers the production of functional TGF-β1, leading to a TGF-β-driven autocrine loop facilitating the effects of the BRAF^V600E^ oncoprotein. This includes the decreased expression of NIS [40] and the promotion of cell migration, invasiveness, and EMT [41]. Tumor-associated-macrophages (TAM) that densely infiltrate BRAF^V600E^-Papillary thyroid cancers also contribute to TGF-β1 in microenvironment [41].

We previously labeled NOX4 as a novel key effector of TGF-β1 in BRAF^V600E^-induced thyroid tumors [7]. The pharmacologic inhibition of the TGF-beta pathway, along with the MAPK pathway, parallels the results obtained from combining NOX4 or p22^phox^ knockdown with MAPK pathway inhibition. Until now the precise mechanism underlying the additive effect of SMAD and MEK inhibition on iodide uptake remained unclear. We demonstrate that TGF-β1-induced signaling through SMAD may especially contribute to the suppression of NIS by altering the recruitment of PAX8 and NKX2.1 *via* NOX4-dependent oxidative damage. Further exploration of the translational implications of these findings in clinical trials will employ a combination of drugs targeting the MAPK pathway and the TGF-β1 pathway.

In conclusion, our findings reveal that NADPH oxidase NOX4, which has notable expression in BRAF^V600E^-mutated thyroid tumor cells and produces H_2_O_2_ in the nuclear environment, repeatedly generates oxidative DNA damage. This promotes increased retention of epigenetic modifiers, such as DNMT1, at the sites of DNA damage alongside DNA repair proteins including OGG1 and MSH2/MSH6. All these factors contribute to blocking the recruitment of two vital transcriptional factors: PAX8 and NKX2.1, which are involved in the transcription of genes associated with thyroid differentiation. The co-inhibition of the MAPK pathway by combining Dabrafenib/Trametinib (which diminishes the expression levels of MSH2/MSH6 and DNMT1) with the TGF-β1 pathway by EW7197 (which reduces NOX4 levels), enhances the recruitment of the two transcription factors to the chromatin (Fig. 10).

**Figure 10:**
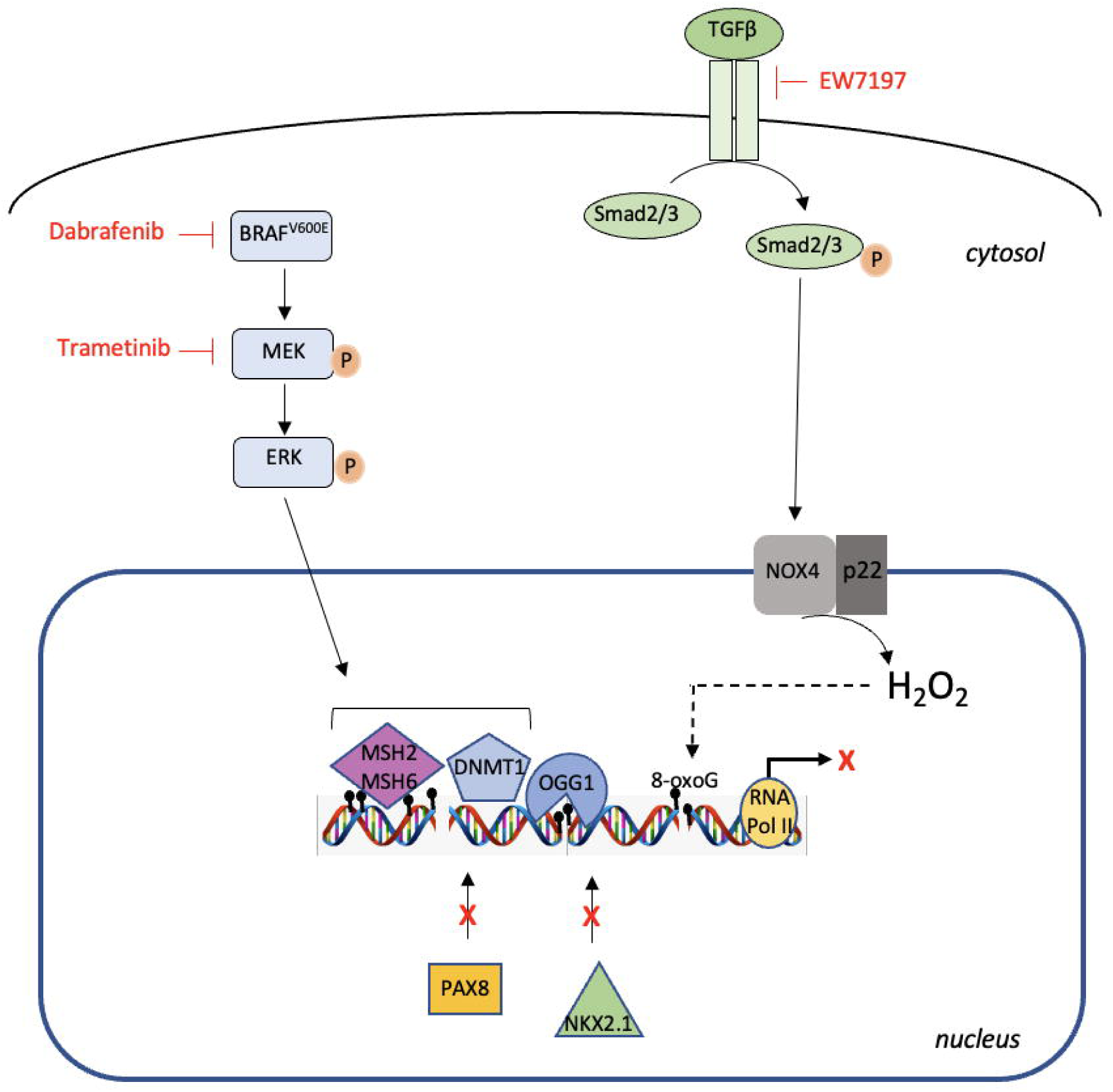
Model proposed for the mechanism. The NADPH oxidase NOX4, which is well expressed in BRAF^V600E^-mutated thyroid tumor cells and produces H_2_O_2_ in the nuclear environment, generates oxidative DNA damage, promoting longer retention of epigenetic modifiers, such as DNMT1, at sites of DNA damage via the interaction with DNA repair proteins including OGG1 and MSH2/MSH6. All contribute to preventing the recruitment of the two key transcription factors: PAX8 and NKX2.1, involved in the transcription of genes involved in thyroid differentiation. Co-inhibition of the MAPK pathway by the combination of dabrafenib/trametinib that decreases MSH2/MSH6 and DNMT1 expression levels and the TGF-β1 pathway by EW7197 that decreases NOX4 levels potentializes the recruitment of PAX8 and NKX2.1 to the chromatin.

## Methods

### Cell culture

The cell lines BCPAP and 8505C were procured from the DSMZ-German Collection and were maintained at a temperature of 37°C and a CO_2_ level of 5% in DMEM (Dulbecco’s Modified Eagle Medium) with 4.5 g/L glucose (Life Technologies), and RPMI-1640 (Life Technologies), respectively. Both mediums were supplemented with 10% (vol/vol) FCS (Life Technologies) and 100 mg/ml solution of penicillin/streptomycin (Life Technologies). All cell lines underwent the authentication procedure STR analysis according to the global standard ANSI/ATCC ASN-0002.1-2021 (2021). Primary human thyroid cells were cultured following the method described previously [42].

### PDX generation and PDX cell line

The PDX563 cell line was derived from a 76-year-old female with BRAF^V600E^-mutated anaplastic thyroid cancer (ATC). She was initially diagnosed with a 3-cm right papillary thyroid carcinoma with extrathyroidal extension in 2010. Treatment involved a total thyroidectomy followed by the administration of 100 mCi I-131. Post-treatment, the control I-131 whole-body scintigraphy and FDG-PET presented as normal. Under LT4 treatment, her serum thyroglobulin level was 0.8 ng/mL. However, in 2017, the serum thyroglobulin level elevated to 3.3 ng/mL. A CT scan revealed a 13-mm right latero-tracheal mass, a 4-mm lymph node in the left compartment VI, and pulmonary micro-nodules. FDG-PET indicated high uptake in the latero-tracheal mass, and a biopsy confirmed a recurrence of the papillary carcinoma. Following a second treatment with 100 mCi of I-131, no uptake was detected on post-therapy whole-body scintigraphy. By the end of 2018, FDG-PET displayed increased uptake in all lesions and a new uptake in the right arm. A biopsy of the right arm revealed an anaplastic carcinoma of the thyroid with significant macrophage infiltration. The patient was treated with a dabrafenib-trametinib combination and external beam radiation to the neck and the right arm. Unfortunately, despite treatment efforts, the neck tumor continued to grow, leading to the patient’s death in 2021.

Fresh tissue biopsy fragments of ATC were implanted into the sub-renal capsule of NOD scid gamma (NSG) mice, acquired from Charles River Laboratories. Xenografts were subsequently propagated subcutaneously from mouse to mouse up to five passages to create a viable tumor bank. A patient-derived cell line was developed from PDX samples. These samples were processed through enzymatic digestion using a tumor dissociation kit (Miltenyi Biotec, #130-095-929,) and mechanical degradation using the gentleMACs^TM^ dissociator. The cells were cultivated using DMEM/F-12+GlutamMAX^TM^ 10% FBS solution (Life Technologies) and 10% enriched solution with 0.4 μg/ml hydrocortisone (Sigma, #H4001), 8.4 ng/ml cholera toxin (Sigma, #C9903), 24 μg/ml adenine (Sigma, #A2786), and 5 μM ROCK inhibitor (Sigma, #Y0503), until a stable proliferation of tumor cells was observed.

### Ethics

The patient participating in the study was fully informed and signed an informed consent form. The MATCH-R trial received approval from the ethics committee at Institut Gustave Roussy, as well as the French National Agency for Medicines and Health Products Safety (ANSM), and is conducted following the Declaration of Helsinki. All animal procedures and studies received approval from the French Ministry of Higher Education, Research, and Innovation (APAFIS#2790-2015112015055793). The MERAIODE study is an investigator-initiated trial sponsored by Gustave Roussy, conducted within the French Endocan-TuThyRef network in approved by ethics committees and national authorities and in accordance with the Declaration of Helsinki. All patients gave their written informed consent

### Cell treatments

For N-Acetyl-L-cysteine (NAC) (Sigma, #A7250) and Diphenylene iodonium (DPI) (Sigma, #300260) treatments, cells were exposed to indicated doses and times at 37°c in their medium and washed once with PBS prior to harvesting. For dabrafenib plus trametinib treatment, BCPAP cells were exposed to 100 nM dabrafenib (Selleck, #S2807) and 25 nM trametinib (Selleck, #S2673) for indicated time at 37°C in DMEM with FCS, 8505C cells were exposed to 100 nM dabrafenib and 5 nM trametinib to indicated time at 37°C in RPMI with FCS and PDX cells were exposed to 100 nM dabrafenib and 5 nM trametinib to indicated time at 37°C in DMEM/F12 with FCS. After treatments, cells were washed once with PBS prior to harvesting. For Vactosertib (EW-7197) (Selleck, #S7530) treatment, cells were exposed to 1 µM of EW-7197 for indicated time at 37°C in their respective medium and washed once with PBS prior to harvesting.

### Tight chromatin fractionation and whole-cell protein extraction

The tight chromatin fractionation protocol was adapted with slight modifications from O’Hagan et al [22]. Briefly, cell pellets were suspended in buffer A (10 mM HEPES pH 7.9, 10 mM KCl, 1.5 mM MgCl2, 0.34 M sucrose, 10 % glycerol, 1mM DTT, 1 × protease and phosphatase inhibitor cocktail (Sigma, #04693116001 and #04906845001). Triton X-100 was then added to the cell suspension to reach a final concentration of 0.1% and incubated for 5 min on ice before centrifugation for 4 min at 1300 × g, 4°C. The supernatant, enriched in cytoplasm soluble proteins, was discarded and the nuclei pellet was washed with buffer A. The pellet was then resuspended in buffer B (3 mM EDTA, 0.2 mM EGTA, DTT 1 mM, 1 × protease, and phosphatase inhibitor cocktail) and incubated for 10 min on ice. After that, the nuclei suspension was centrifuged for 4 min at 1700 × g, 4°C. The supernatant was discarded and the pellet, representing the chromatin fraction, was washed in buffer B and centrifuged for 1 min at 10,000 g, 4°C. The chromatin fraction was washed with buffer C (50 mM Tris/HCl pH 8, NaCl 0.45 M, IGEPAL 0.05 %, 1 × protease, and phosphatase inhibitor cocktail) to solubilize proteins weakly bound to chromatin and then centrifuged for 1 min at 10,000 g, 4°C. The remaining pellet was lysed in TEX buffer (100 mM Tris·HCl pH 7.0 containing 2.5% (wt/vol) SDS, 1 mM EDTA, 1 mM EGTA, 4 M urea, and a mixture of phosphatase and protease inhibitors using Qiashredder (Qiagen, #79654) and referred to as tight chromatin. Whole-cell extracts were prepared from 1/10 of the pellet collected after treatment before beginning the tight chromatin isolation. Vinculin and Lamin B immunoblotting served as cytoplasmic and nuclear controls, respectively.

### siRNA knockdown

Cells were transfected at 50–60% confluence with specific human siRNA against NOX4 (Thermo Scientific, silencer select siRNA, #4392421, s224160), scrambled siRNA control (Thermo Scientific, #4392420 and Horizon Perkin Elmer company, On target plus non-targeting control pool, #D-001810-10-05), siRNA against p22^phox^ (Horizon Perkin Elmer company, on target-plus human CYBA siRNA, #L-011020-02-0005), siRNA against OGG1#1 (Thermo Scientific silencer select siRNA, #4390824, s9836), siRNA against OGG1#2 (Thermo Scientific silencer select siRNA, #4390824, s9837), siRNA against DNMT1 (Horizon Perkin Elmer company on target-plus siRNA pool, #L-004605-00-0005), siRNA against MSH2 (Thermo Scientific, Stealth Select RNAiTM, #1299001), and siRNA against MSH6 (Eurogentec, Target sequences: 5′-CCCUGGCAAACAGAUUAAA-3′, [13]) using the INTERFERin transfection reagent (Polyplus-Transfection, # POL101000036) following the manufacturer’s protocol.

### Protein extraction and Western blotting

The Western blot analysis was conducted using lysates prepared as described previously [43]. The membranes were probed with primary antibodies: anti-NOX4 (Abcam, #ab109225, RRID:AB_10861375); anti-p22^phox^ (Santa Cruz Biotechnology, #Sc-130551, RRID:AB_2245805); anti-LaminB1 (Abcam, #ab133741, RRID:AB_2616597); anti-MSH6 (Protein Tech, #A3000 22 A); anti-MSH2 (Abcam, #ab70270); anti-OGG1/2 (G5) (Santa Cruz Biotechnology, #Sc376935); anti-Vinculin (Abcam, #ab130007); anti-DNMT1 (Abcam, #ab13537); anti-DNMT3a (Abcam, #ab2850); anti-DNMT3b (Cell signaling, #695202); anti-PAX8 (Cell signaling, #59019); anti-TTF1 (NKX2.1) (Cell signaling, #123735), anti phospho-p44/42 ERK (Cell signaling, #4370); anti-ERK (Cell signaling, #4696); anti phosphor-MEK1/2 (Cell signaling, #9121); anti-MEK (Cell signaling, #4694); anti-BRAF^V600E^ (Spring, #E19290); anti-phospho-Smad3 (Abcam, #ab515663); anti-Smad3 (Thermo Scientific, #MA515663) or anti-Myc-Tag (Cell signaling, #22765) overnight at 4°C with constant agitation. They were then washed three times using TBS-T and treated with either goat anti-rabbit IgG-HRP antibody (Southern Biotech RRID:AB_2687483) or goat anti-mouse IgG-HRP antibody (Agilent, #P0447, RRID:AB_2617137) for 45 min at room temperature. Following another series of washes – three times with TBS-T – the proteins were visualized through enhanced chemiluminescence (Thermoscientific, #34577).

### Flow cytometry analysis of 8-oxoG

Cells were fixed in 70% ethanol at -20°C and washed with PBS before being permeabilized with 0.1% Triton X-100/PBS for 15 min at room temperature. They were then washed with PBS and blocked in 2.5% bovine serum albumin/PBS for 30 min at room temperature. The cells were treated with 50µg/ml RNAse and incubated with anti-8-oxodG (Biotechne/R&D systems #4354-MC-050, 1:50 diluted in 2.5% BSA/PBS) for 1 h at 37°C. The cells were then washed once with 0.2% BSA/PBS and incubated with the secondary antibodies Alexa 488 anti-mouse (Invitrogen, #A11017, RRID:AB_2534084) in 0.2% BSA/PBS for 1 h at 37°C before propidium iodide staining. Cytofluorimetric acquisition and analysis were conducted on a BD Accuri C6 plus Flow cytometer.

### Nuclear H_2_O_2_ production by FACS analysis

The measurement of nuclear H_2_O_2_ was accomplished using NucPE1 (Nuclear Peroxy Emerald 1) (Med Chem Express, #1404091-23-1) [44]. The cells were incubated with 10 µM NucPE1 for 45 min in the dark. After incubation, the cells were washed and analyzed by flow cytometry.

### Plasmid transfection

BRAF^V600E^ Construct—cDNA was synthesized with Maxima reverse transcriptase (Thermo Scientific, # EP0741) by oligo(dT) priming of total RNA from BCPAP cells. The BRAF^V600E^ ORF was amplified using Phusion Plus DNA polymerase (Thermo Scientific, # F630L) and cloned into pcDNA™3.1/V5-His TOPO^®^ vector (Thermo Scientific, # K480001) following the manufacturer’s instructions. Primer sequences used for BRAF^V600E^ amplification were: forward: 5’-CACCATGGCGGCGCTGAGCGGTG-3’ and reverse: 5’ - TCAGTGGACAGGAAACGCACCATATCC-3’. The construct was verified by sequencing.

Twenty-four hours before the experiment, human thyrocytes from the primary culture (2.5 × 10^5^ cells/well) were seeded in 6-well plates. Cells were transiently transfected following the manufacturer’s instructions using 3 μl of X-tremGENE HP DNA transfection reagent (Sigma, #6366236001) and 1 μg of total DNA, which consisted of the BRAF^V600E^/pcDNA3.1 plasmid. Experiments were conducted 48 h post-transfection.

### Plasmid/siRNA cotransfection

Twenty-four hours before the experiment, BCPAP and PDX cell lines (4 × 10^5^ cells/well) were seeded in 6-well plates. Cells were transiently transfected according to the manufacturer’s instructions using 4 μl of jetPRIME siRNA and DNA transfection reagent (Polyplus-Transfection, #101000046), 10 µM of siRNA, and 400 ng of total DNA (Myc-OGG1/pcDNA3.1 plasmid [45], kindly provided by D. Sidransky (Addgene plasmid 18709). Experiments took place 72 h post-transfection

### Genomic DNA extraction

Genomic DNA was extracted using the DNeasy Blood & Tissue kit (Qiagen, #69504). All extraction steps were performed under low-light conditions. Additionally, 50 μM N-tert-butyl-α-phenylnitrone (stock solution: 28 mM in H2O; Sigma, #B7263,) was added to each DNeasy Blood & Tissue buffer to protect the DNA from further oxidation.

### RNA extraction, cDNA synthesis, and real-time PCR

Total RNA from cell samples was purified with the Nucleospin RNA II kit (Machery Nagel, #740955-50). RNA yield was determined using a spectrophotometer (NanoDrop Technologies, Wilmington, USA). DNase-treated RNA (1 µg) was reverse-transcribed using Maxima reverse transcriptase (Thermo Fisher Scientific, #EP0743) and oligo-dT in a total reaction volume of 20 μl of PCR buffer, adhering to the manufacturer’s protocol for 60 min at 55°C. Quantitative PCR (qPCR) was performed on an ABI 7500 system (Applied Biosystems) using Taqman gene expression assays (Thermo Fisher Scientific): SLC5A5 (Hs00950365_m1); NOX4 (Hs04980925_m1) and TSHR (Hs01053846_m1)

### Tissue samples and Immunohistochemical study

Twenty one patients with metastatic radioactive iodine refractory BRAF^V600E^-mutated differentiated thyroid cancer were included in a phase II redifferentiation trial with dabrafenib-trametininb and ^131^I. The study design, the characteristics of the patients and the trial results were published [21]. Fifteen tumors from these patients were analyzed by immunohistochemistry (Supplemental Table 1). The immunohistochemical (IHC) study was performed on 3-μm-thick deparaffinized slides obtained from paraffin blocks on Ventana Automated Immunostainer (BenchMarker Ultra, Ventana Medical System, Inc,) according to the manufacturer’s manual. All the technical part was handled by the experimental and translational pathology platform at Gustave Roussy (PETRA). Primary monoclonal (M) and polyclonal (P) antibodies were directed to: NOX4 (home-made antibody mAb8E9 [46]; 1/500), MSH2 (Millipore ; clone FE11 ; 1/20); MSH6 (BD Bioscience ; clone 44 ; 1/400) ; OGG1 (antibodies online ; 6940242 ; 1/200) ; p-SMAD3 (ABCAM ; ab51451 ; 1/500) ; PAX8 (Diagomics ; C-P008 ; prediluted) and NKX2.1 (Diagomics ; 8G7G3/1 ; prediluted). The specificity of the reactions was verified by appropriate positive and negative controls for each antibody.

IHC slides were scored by a pathologist using a standard light microscope. IHC slides with significant tissue artifacts in the tumor area that would not allow IHC assessment were excluded from analysis. All scores were performed at 20X magnification. Each markers expression scoring was assessed in malignant cells using a standard microscope and a semi-quantitative approach combining both the percentage of positive cells (0-100) and the intensity of tumor immunostaining (0, no staining; 1+, for weak, 2+, for moderate; and 3+, for strong expression), and then an H-score was calculated using the following formula: H-score = [1 × (% 1+ cells) + 2 × (% 2+ cells) + 3 × (% 3+ cells)]. Any immunoreactivity with an intensity of less than 1+ was considered background or nonspecific (“0”). The H-score range from 0 (no tumor staining) to 300 (strong staining of all tumor cells analyzed).

Statistical methods : Normality of H-score between the markers was assessed with the Shapiro-Wilk tests. Spearman’s coefficient was used to assess the correlation between each H-score . Correlation was judged very strong from 1 to 0.9, strong from 0.9 to 0.7, moderate from 0.7 to 0.5, low from 0.5 to 0.3 and poor from 0.3 to 0. The alpha risk was set to 0.05.

### ChIP-qPCR assays

ChIP assays were carried out on BCPAP cells using a ChIP-IT Express kit (Active Motif, #53008), following the manufacturer’s instructions. Briefly, the cross-linked cell chromatin was sheared through sonication for 15 min at 40 W (duty factor: 20%, peak incident power:200, cycles per burst: 200) with a Covaris S220 (Woodingdean, UK). Subsequently, the chromatin fragments were immunoprecipitated with 1 µg of anti-NKX2.1 (TTF1) antibody (Cell Signaling Technology, #12373, RRID:AB_2797895) or an equal amount of rabbit IgG isotype control (Agilent, #0903). The DNA bound to the chromatin immunoprecipitates was eluted and analyzed using qPCR with FastStart Universal SYBR Green Master (ROX) (Sigma/MERK, #4913850001) for detecting the TG proximal promoter sequence with specific primers (TG forward: GAGTAGACACAGGTGGAGGGA and TG reverse: GCTTTTATAGAGCTGCCGTTGG). The fold enrichment of the ChIP samples, relative to the IgG samples, was calculated via the slope of a standard curve established by performing qPCR with the primer set on known DNA quantities of input DNA (Active Motif).

### **γ**H2AX ChIP-seq analysis

Enriched DNA from ChIP and Input DNA fragments were end-repaired, extended with an ‘A’ base on the 3′end, ligated with indexed paired-end adaptors (NEXTflex, Bioo Scientific) using the Bravo Platform (Agilent), size-selected after 4 cycles of PCR with AMPure XP beads (Beckman Coulter) and amplified by PCR for 10 cycles more. The final libraries were purified, pooled together in equal concentrations and subjected to paired-end sequencing (100 cycles: 2x50) on Novaseq-6000 sequencer (Illumina) at Gustave Roussy.

Quality control of reads was performed by FastQC (RRID:SCR_014583) [47], then good reads were mapped with BWA (RRID:SCR_010910) [48] to the hg38 reference from UCSC and the PCR duplicates were identified by SAMtools (RRID:SCR_002105) [49]. Discared reads were those not marked as primary alignments, were unmapped, were mapped to multiple locations, and were not paired. Those evaluation and filtering were carried out by SAMtools [49]. The broad peaks were called by MACS2 [50] with a broad-cutoff parameter set to 0.05. Homer [51] annotation of peaks was performed to evaluate GC bias between samples, and PCA and correaltion Heatmap were made by deepTools [52]. The consensus peaksets by condition were identified by BEDTools (RRID:SCR_006646) [53]. Reads marked as duplicates or were mapped to blacklisted regions, were removed, then signal enrichment was scaled to 1 million mapped reads, then normalized to input sample by deepTools [52]. To merge signal by condition, WiggleTools [54] was used to create a mean merged Bedgraph by condition, then convert it into BigWig file with the UCSC bedGraphToBigWig [55] function. Next, ChIPseeker [56] R package was used to plot the consensus peaks coverage along chromosomes (under R 4.3.1). All computations used conda [57] environments, making results highly reproducible.

### ATAC sequencing analysis

For ATAC-sequencing, cells from BCPAP, 8505C, and PDX were extracted and the library preparation was conducted according to the instructions provided in the ATAC-Seq Kit (Active Motif, #53150). The quality of raw reads was assessed using FastQC [47] and FastQScreen [58], indicating good overall quality for ATACSeq. Graft reads from ATAC-Seq was mapped onto the human genome from Ensembl, version GRCh38.104. We used Sambamba [59] to remove reads with mapping quality below 30, as well as any reads without mates mapped in a proper pair. Although duplicate reads were marked with Sambamba, they were not eliminated for downstream analysis. We shifted ATAC-Seq reads using DeepTools [60], as proposed in the protocol by Buenrostro et al. [61]. Additional quality controls for ATAC-Seq were performed with ATACSeqQC [62]. Since the reads were previously shifted, ATAC-Seq was called using the same parameters as other libraries. We annotated peaks using Homer [51] and conducted quality controls for peak calling with DeepTools, ChipSeeker [63], and Python scripts developed in-house. The differential peak analysis was performed using edgeR [64] along with CSAW [65].

### Long read library preparation and sequencing

Genomic DNA was quantified using a Qubit Fluorometer (Thermo Fisher) and size average was determined using Fragment Analyzer (Agilent). Sequencing libraries were prepared using Ligation Sequencing Kit V14 (SQK-LSK114, ONT, Oxford, UK). Briefly, 1000ng of gDNA were repaired and Ultra II end-prep according to the manufacturer recommendations. After elution, quantity was evaluated using Qubit fluorometer and adapter was ligated. Only flow cells with at least 5,000 pores at the initial scan were used for sequencing. After AMPure clean-up, 15 fmol of prepared library was loaded onto R10.4.1 flow cells (FLO-PRO114M, ONT) on a PromethION P-48 device (ONT) at Gustave Roussy with MinKNOW and super-accuracy base-calling mode selected with m5C and hm5C analysis.

### DNA methylation analysis

Data were sequenced on flowcell R10.4 on a PromethION 48 using Minknow (v5.5.3). Output pod5 files were base called and methylations (5mCG and 5hmCG) were detected on Dorado basecaller (v0.5.3) with, respectively, the models dna_r10.4.1_e8.2_400bps_sup@v4.2.0 and “dna_r10.4.1_e8.2_400bps_sup@v4.2.0_5mC_5hmC@v1”, and the options “--emit-moves -- no-trim -b 1728”. With a homemade python3 script using pysam, the reads, with mean quality score (“qs”) under 10 and length under 200, were removed. The BAM were aligned on Grch38 (build 109 from Ensembl), customized with the Lambda phage sequence (NC_001416.1 from NCBI), using Dorado aligner (v0.5.3) and the options “--secondary = no” and “--bandwidth 500,20000”. The aligned BAM was concatenated in a single BAM, sorted, indexed, and then split by chromosome using Samtools (v1.11) [66] sort, index, and view in this order. For each BAM, the 5mCG and 5hmCG were separated using modkit (v0.2.6) adjust-mods with “--ignore h” (ignore 5hmCG) or “--ignore m” (ignore 5mCG). Then, methylations were extracted in a bed file with “modkit pileup” and the option “--cpg”. A differential methylated regions analysis on transcript features (exons, introns, TSS, promoters) was performed using R (v4.1.2) package GeneDMRs (v1.1.0) [67] with the reference from the UCSC (Clade:Mammal, Genome:human, Assembly:Dec.2013(GRCh38/hg38), Group:Genes and Gene Predictions, Track: GENECODE V46, Table: knowngene). The heatmap were produced with pheatmap (v1.0.12) [68], the volcanoplot with EnhancedVolcano (v1.12.0) [69], and the piecharts with ggplot2 (v3.4.4) (RRID:SCR_014601) [70]. The IGV snapshots were obtained on IGV (v11.0.21).

### Quantification and statistical analysis

Statistical analyses were performed using GraphPad Prism software. Data were analyzed by One- or Two-way ANOVA or Student’s t-test, with the minimum level of significance set at P<0.05.

## Acknowledgments

The authors would like to thank Pr Martin Schlumberger and Dr Patricia Kannouche for their critical reading of the manuscript. They would like to thank Pr Isabelle Borget for her statistical analysis of IHC data. They would like to thank the SIRIC EpiCure (INCa-DGOS-Inserm-ITMO Cancer_18002) for its support. They are also indebted to Audrey Naimo from the genomic platform (NSERM US23, CNRS UMS 3655, AMMICa, Gustave Roussy) for her assistance with sequencing.

## Author contributions

M. Radom performed experiments and conducted data/statistical analysis; C. Buffet performed experiments and conducted data/statistical analysis; J. Cazarin, M. Harinquet and C. Coelho de Faria initiated experiments; F. Brayé and C. Nobre contributed to the preparation of the PDX cell line; M Aglave, Y Mesloub and T. Dayris conducted bioinformatic analysis; N. Droin supervised sequencing experiments, K. Godefroy performed Immunohistochemistry (IHC), M-A Bani performed IHC analysis, A Al Ghuzlan analysed and selected human samples for thyrocytes preparation; S. Leboulleux provided tumor samples within the MERAIODE trial, L. Lamartina contributed to the development of the project; C. Dupuy designed, supervised experiments, provide funding and wrote the manuscript.

