## Supplementary figures for "NOX4 prevents the recruitment of PAX8 and NKX2.1 to chromatin in BRAF-mutated thyroid cancer cells"

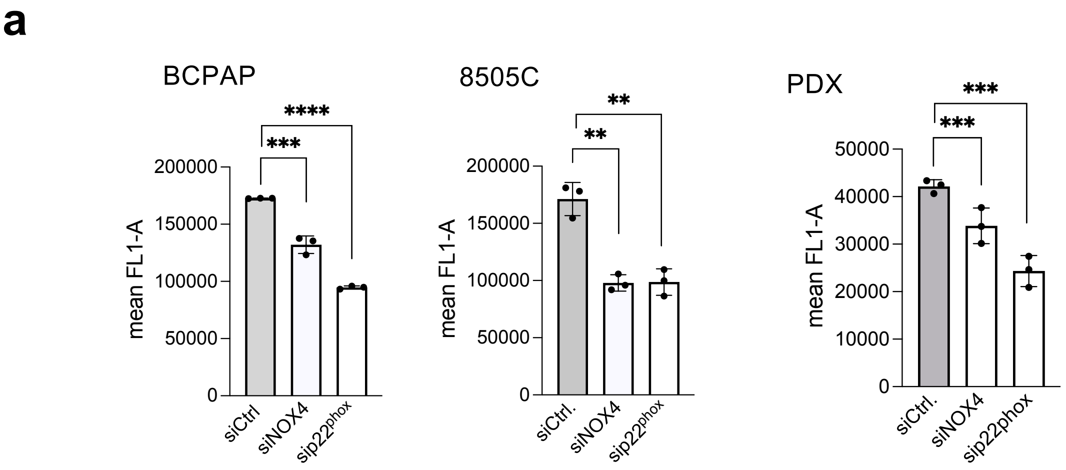


**Supplementary Figure 1:** Knockdown of NOX4 or p22$^{phox}$ reduces oxidative-DNA damage **a**) The nuclear level of ROS was measured in BRAF-mutated thyroid cells by NucPEI fluorescence using flow cytometry. The cells were transduced with siRNA control or siRNA against NOX4 or p22$^{phox}$ for 72 h. Graphs show the quantification of the fluorescence mean. Values are mean ± SE. *$p < 0.05$, **$p < 0.01$, ***$p < 0.001$ and ****$p<0.0001$ ($n = 3$).

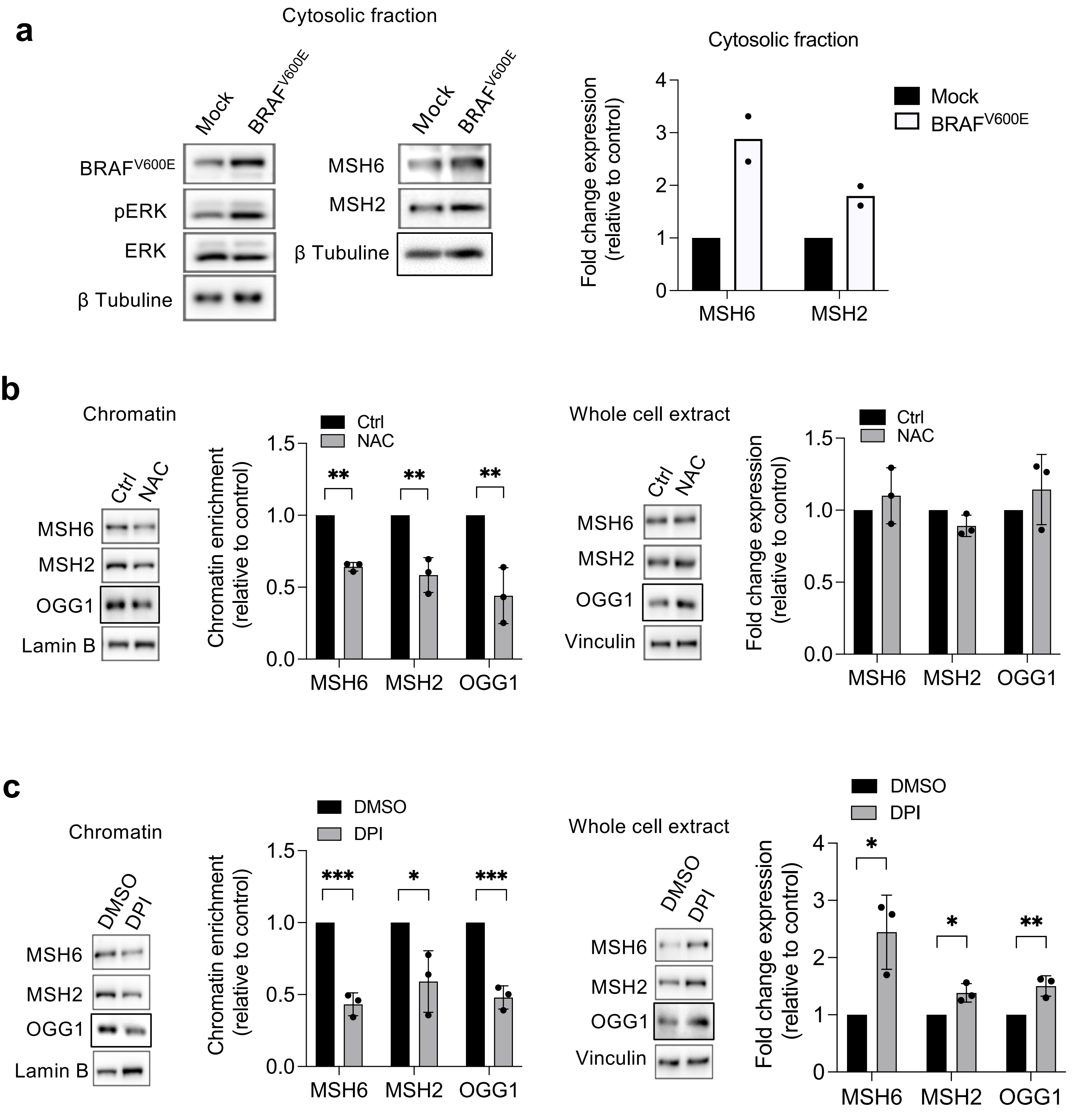


**Supplementary Figure 2**: Anti-oxidant (NAC) or NADPH oxidase inhibitor (DPI) alters the recruitment of OGG1, MSH2, and MSH6 to chromatin in BRAF-mutated thyroid cells. **a**) Human primary thyrocytes were transiently transfected with empty vector (Mock) or BRAFV600E plasmids for 48 h and phospho-ERK, ERK, MSH6, and MSH2 protein expression levels were analyzed by Western-blot in cytosolic fractions (n = 2). **b**) BCPAP cells were pre-treated with 5 mM NAC for 2 h before being analyzed for OGG1, MSH2, and MSH6 protein expressions in chromatin fractions and whole-cell extract. **c**) BCPAP cells were pre-treated with 1 µM DPI for 6 h before being analyzed for OGG1, MSH2, and MSH6 protein expressions in chromatin fractions and whole-cell extracts. Densitometry quantification of protein levels normalized to Lamin B or Vinculin levels and presented as chromatin enrichment or fold change compared with control cells. Values are mean ± SE. *$p < 0.05$, **$p < 0.01$, and ***$p < 0.001$ (n = 3).

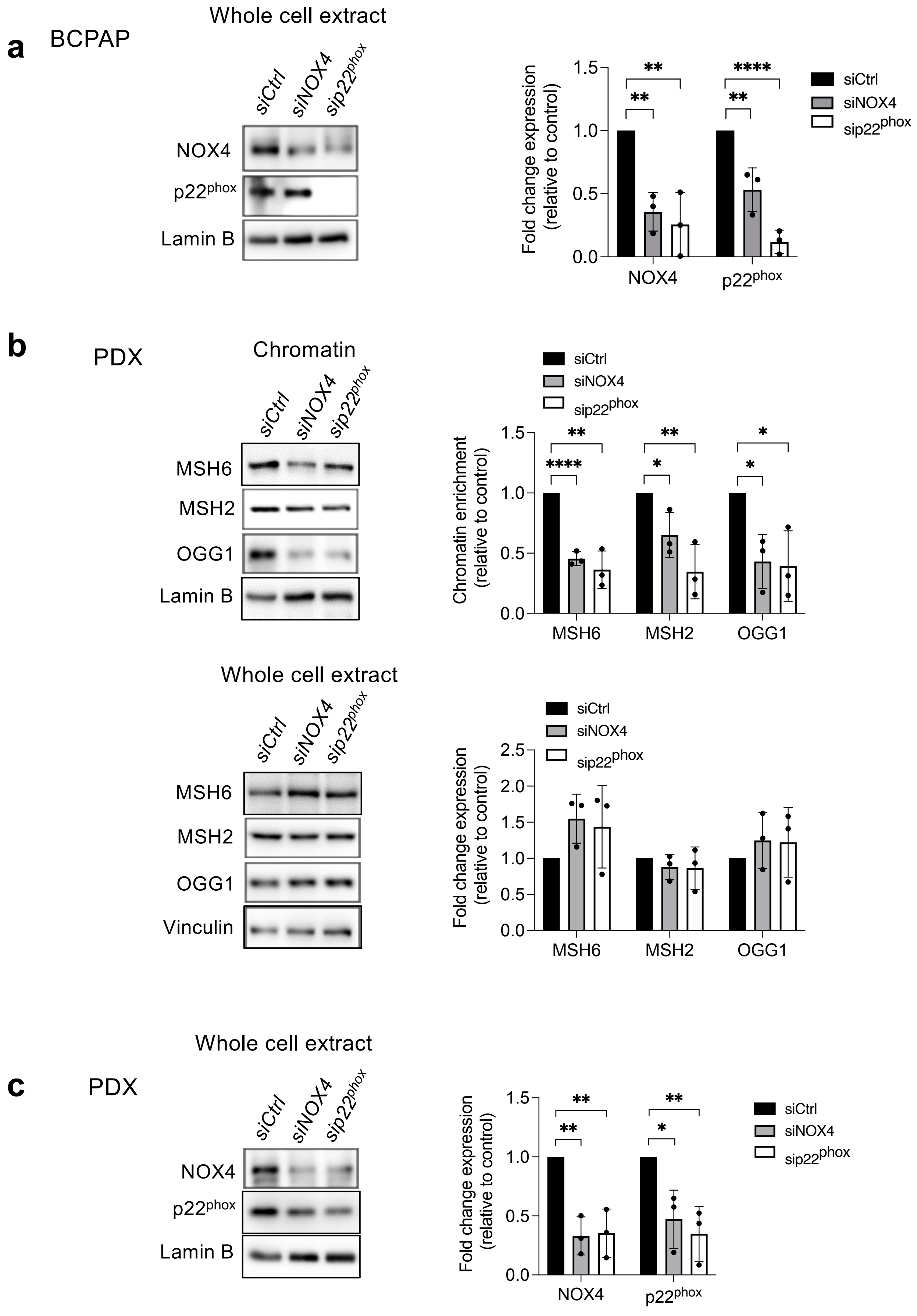


Supplementary Fig. 3

**Supplementary Figure 3** : Knockdown of NOX4 or $p22^{phox}$ reduces the recruitment of OGG1, MSH2, and MSH6 to chromatin in BRAF-mutated thyroid cells. **a**) siRNA knockdown validation of NOX4 and $p22^{phox}$ protein expression levels analyzed by Western-blot in whole-cell extract 72 h after knocking down of NOX4 or $p22^{phox}$ by RNA interference in BCPAP cells. **b**) Western blot analysis of MSH6, MSH2, and OGG1 protein expression levels in chromatin fractions and whole-cell extracts 72 h after knocking down of NOX4 or $p22^{phox}$ by RNA interference in PDX cells. **c**) Western blot analysis of NOX4 and $p22^{phox}$ protein expression levels in whole-cell extract 72 h after knocking down of NOX4 or $p22^{phox}$ by RNA interference in PDX cells. Densitometric quantification of protein levels normalized to loading control levels and presented as chromatin enrichment or fold change compared with siRNA control-transduced cells. Values are mean ± SE. *$p < 0.05$, **$p < 0.01$, ***$p < 0.001$ and ****$p<0.0001$ ($n = 3$).

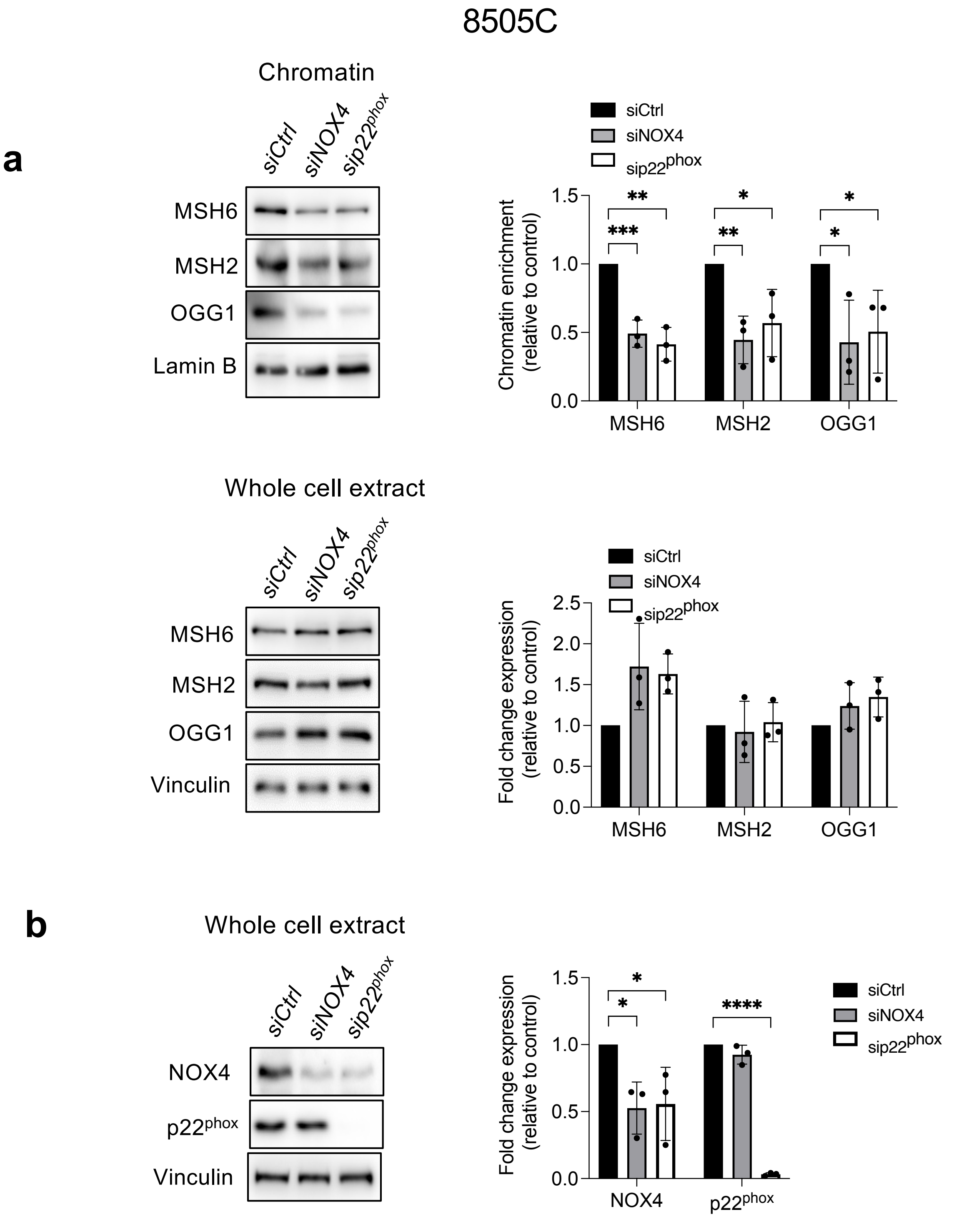


**Supplementary Figure 4**: Knockdown of NOX4 or $p22^{phox}$ reduces the recruitment of OGG1, MSH2, and MSH6 to chromatin in BRAF-mutated thyroid cells. **a**) Western blot analysis of MSH6, MSH2, and OGG1 protein expression levels in chromatin fractions and whole-cell extracts 72 h after knocking down of NOX4 or $p22^{phox}$ by RNA interference in 8505C cells. **b**) Western blot analysis of NOX4 and $p22^{phox}$ protein expression levels in whole-cell extracts 72 h after knocking down of NOX4 or $p22^{phox}$ by RNA interference in 8505C cells. Densitometric quantification of protein levels normalized to loading control levels and presented as chromatin enrichment or fold change compared with siRNA control-transduced cells. Values are mean ± SE. *$p < 0.05$, **$p < 0.01$, ***$p < 0.001$ and ****$p<0.0001$ ($n = 3$).

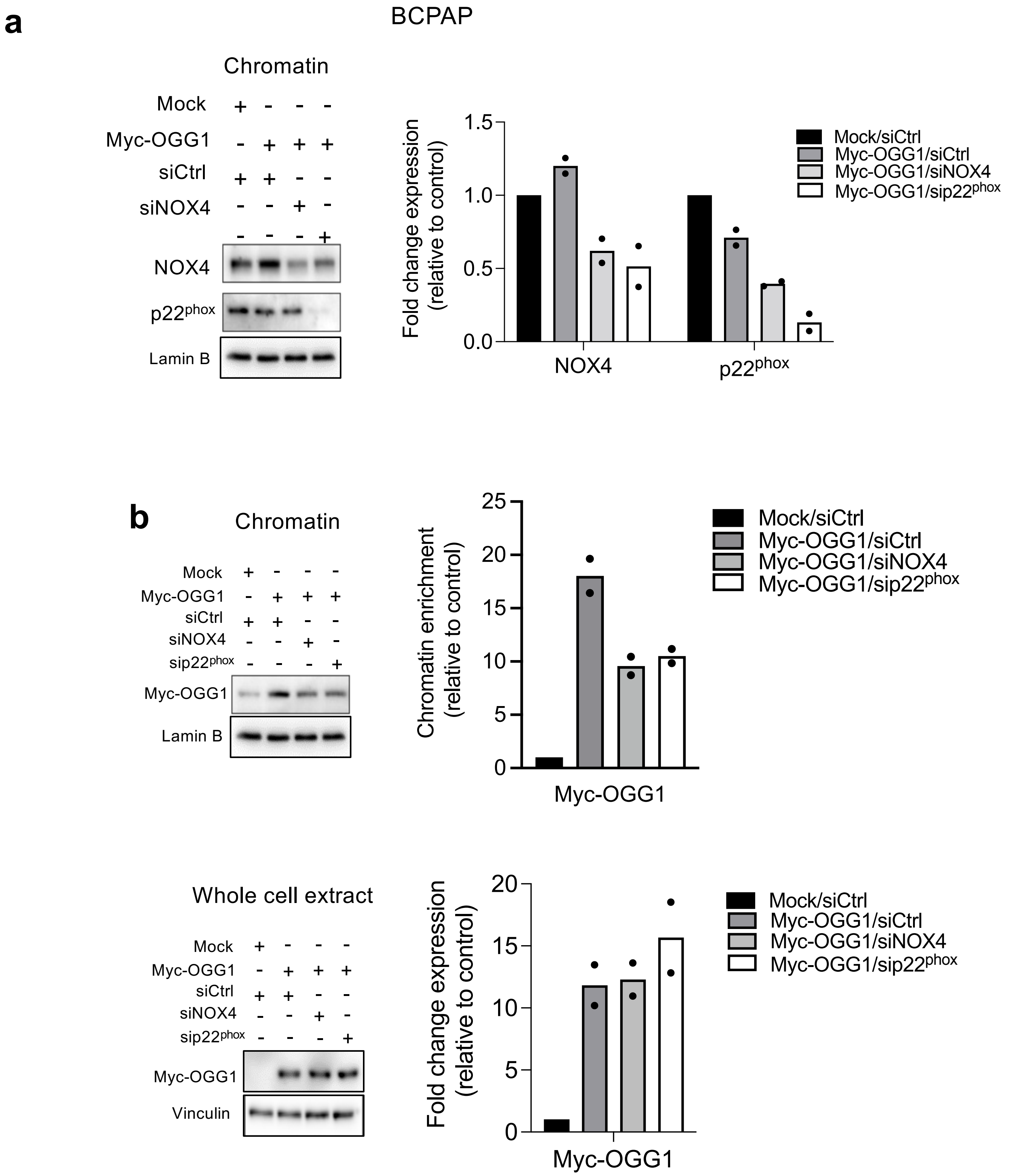


**Supplementary Figure 5.** Knockdown of NOX4 or p22$^{phox}$ reduces the recruitment of OGG1 to chromatin in BCPAP cells. **a, b**) BCPAP cells were transiently transfected with empty vector (Mock) or c-Myc-tagged OGG1 (Myc-OGG1) plasmids in the presence of siRNA control or siRNA NOX4 for 72 h. NOX4, p22$^{phox}$, and Myc-OGG1 protein expression levels were analyzed in chromatin fractions and whole-cell extracts. Densitometry quantification of protein levels normalized to Lamin B or Vinculin levels and presented as chromatin enrichment or fold change compared with control cells (n = 2).

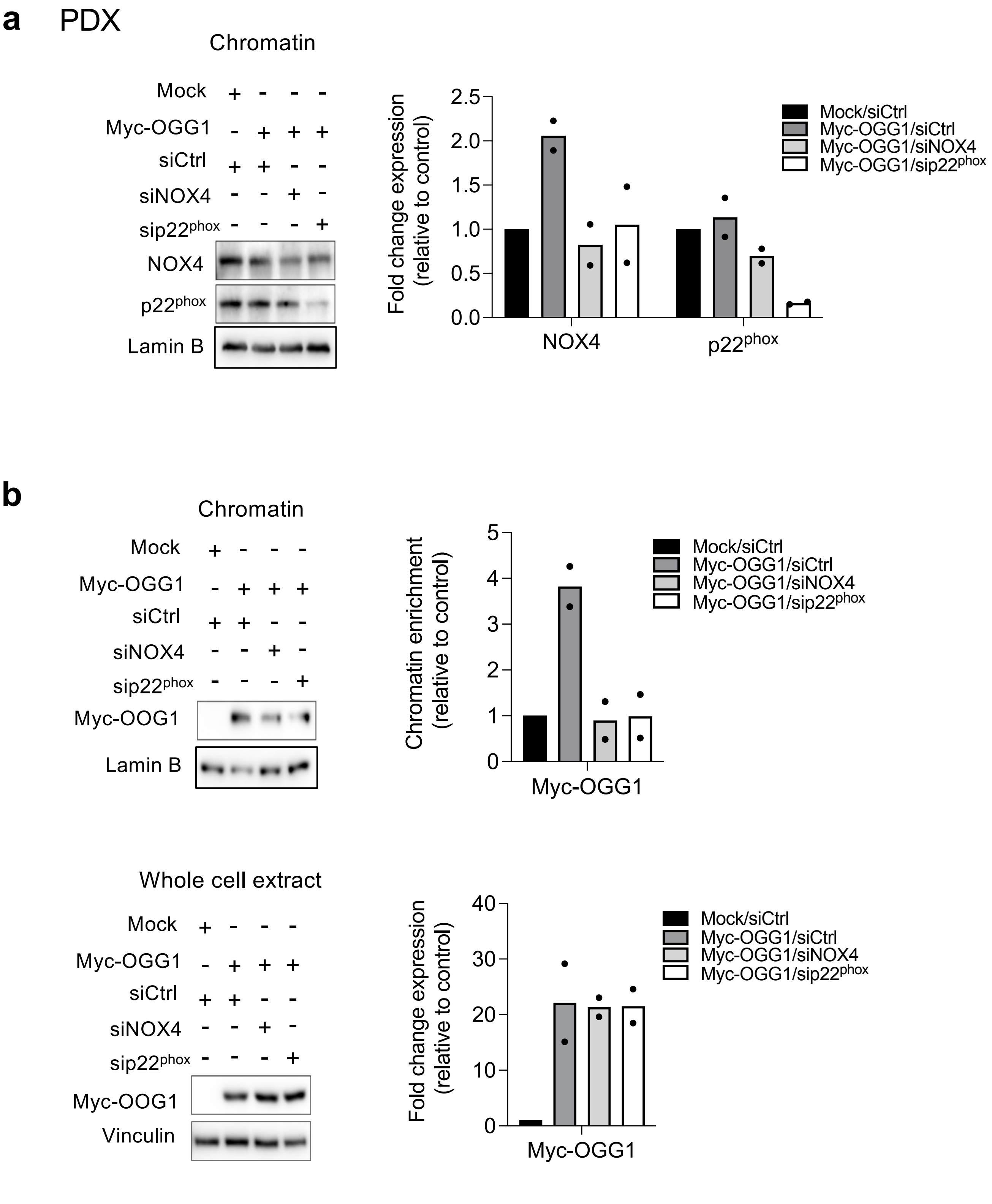


**Supplementary Figure 6**: Knockdown of NOX4 or p22$^{phox}$ reduces the recruitment of OGG1 to chromatin in PDX cells. a**, b**) PDX cells were transiently transfected with empty vector (Mock) or c-Myc-tagged OGG1 (Myc-OGG1) plasmids in the presence of siRNA control or siRNA NOX4 for 72 h. NOX4, p22$^{phox}$, and Myc-OGG1 protein expression levels were analyzed in chromatin fractions and whole-cell extracts. Densitometry quantification of protein levels normalized to Lamin B or Vinculin levels and presented as chromatin enrichment or fold change compared with control cells (n = 2).

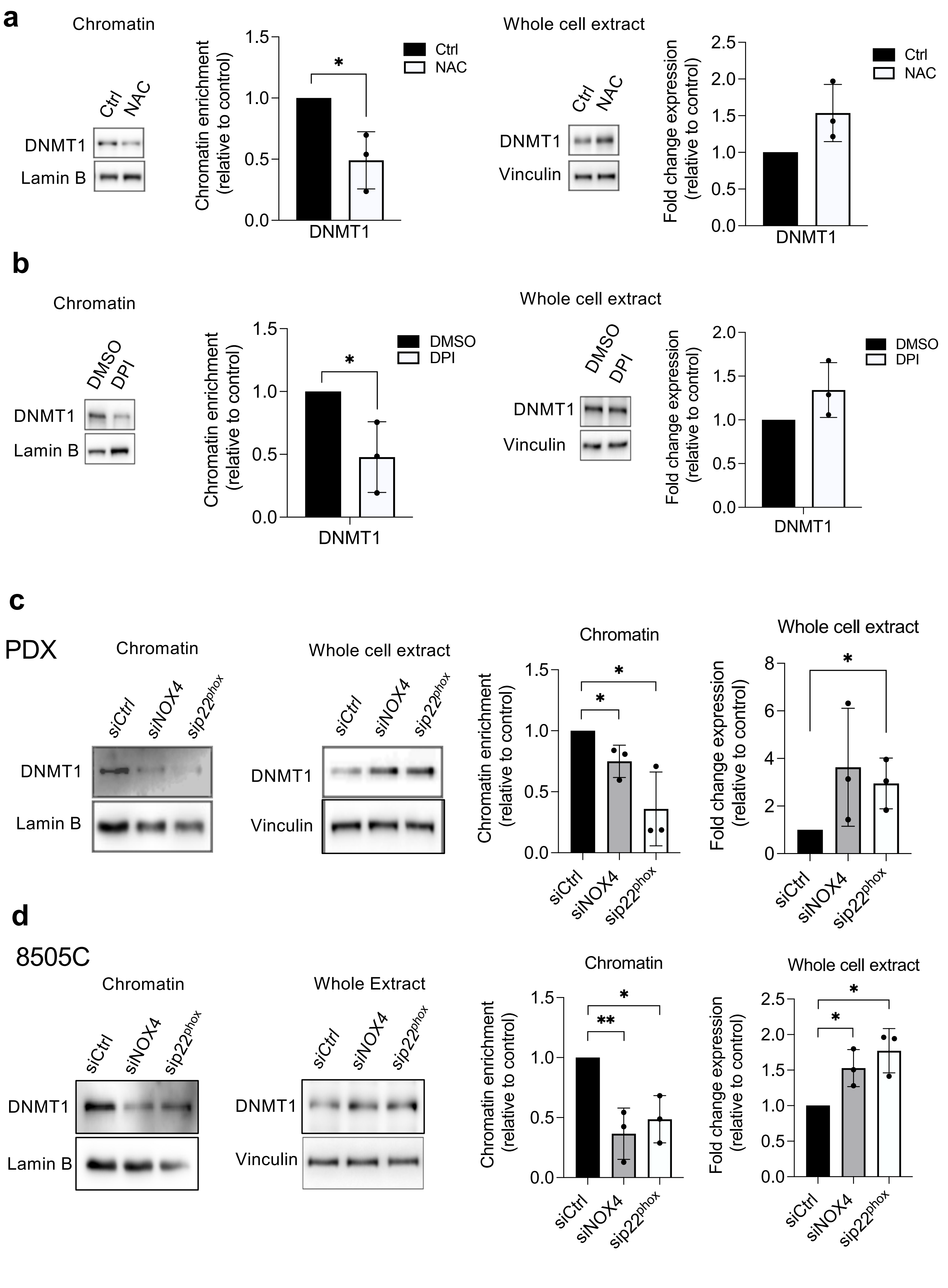


Supplementary Fig. 7

**Supplementary Figure 7**: Anti-oxidant (NAC) or NADPH oxidase inhibitor (DPI) alters the recruitment of DNMT1 to chromatin in BRAF-mutated thyroid cells. **a**) BCPAP cells were pre-treated with 5 mM NAC for 2 h before being analyzed for DNMT1 protein expression in chromatin fractions and whole-cell extracts. **b**) BCPAP cells were pre-treated with 1 μM DPI for 6 h before being analyzed for DNMT1 protein expression in chromatin fractions and whole-cell extracts. **c**) Western blot analysis of DNMT1 protein expression in chromatin fractions and whole-cell extracts 72 h after knocking down of NOX4 or $p22^{phox}$ by RNA interference in PDX cells. **d**) Western blot analysis of DNMT1, the protein expression level in chromatin fractions, and whole-cell extracts 72 h after knocking down of NOX4 or $p22^{phox}$ by RNA interference in 8505C cells. Densitometry quantification of protein levels normalized to Lamin B or Vinculin levels and presented as chromatin enrichment or fold change compared with siRNA control-transduced cells. Values are mean ± SE. $^*p < 0.05$ and $^{**}p < 0.01$ (n = 3).

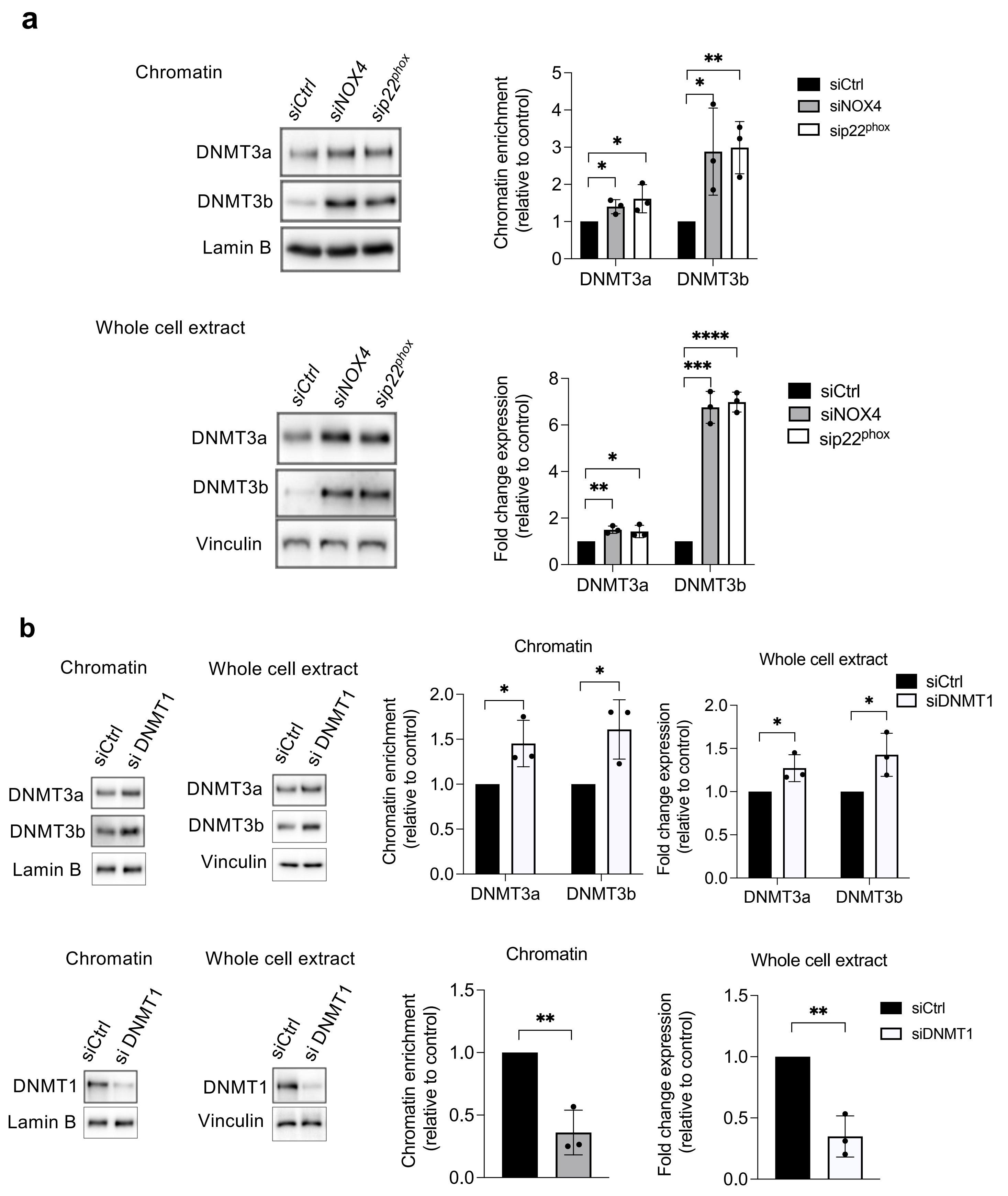


**Supplementary Figure 8**: Knockdown of NOX4 or p22$^{phox}$ increases the recruitment of DNMT3a and DNMT3b to chromatin. **a**) Western blot analysis of DNMT3a and DNMT3b protein expression levels in chromatin fractions and whole-cell extracts 72 h after knocking down of NOX4 or p22$^{phox}$ by RNA interference in BCPAP cells. **b**) Western blot analysis of DNM3a, DNMT3b and DNMT1 protein expression in chromatin fractions and whole-cell extracts 72 h after knocking down of DNMT1 by RNA interference in BCPAP cells. Densitometry quantification of protein levels normalized to Lamin B or Vinculin levels and presented as chromatin enrichment or fold change compared with control cells. Values are mean ± SE. *$p < 0.05$ and **$p < 0.01$ ($n = 3$).

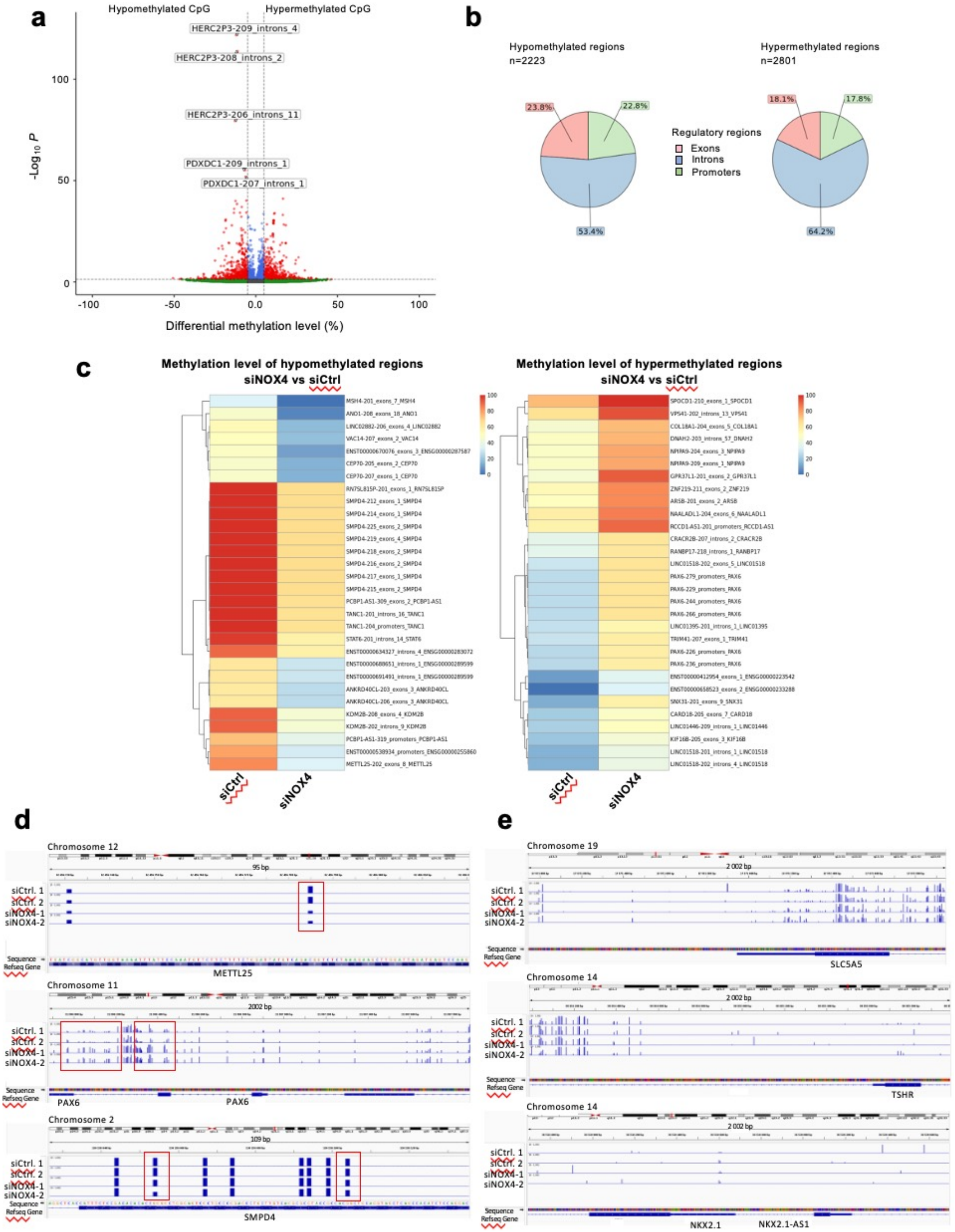


**Supplementary Figure 9**: Comparison of DNA methylation profiles between BCPAP cells transfected with siRNA control and BCPAP cells transfected with siRNA directed against NOX4. **a**) Volcano plots showing the percentage of differentially methylated cytosines for siRNA NOX4 vs siRNA control. Cutoff of adjusted p-value at 0.05 (5%) and differential methylation level at 5%. Green: not significant differentially methylated regions, Blue: significant not differentially methylated regions, and Red: significant differentially methylated regions**. b**) Pie chart showing the distribution of hypermethylated and hypomethylated regions after NOX4 depletion. **c**) Heatmap of Top 30 differentially methylated regions corresponding to annotated genes that are hypomethylated or hypermethylated after NOX4 depletion. **d**) and **e**) IGV representations of the methylation level of some genes**.** The analysis was done from two independent biological replicates.

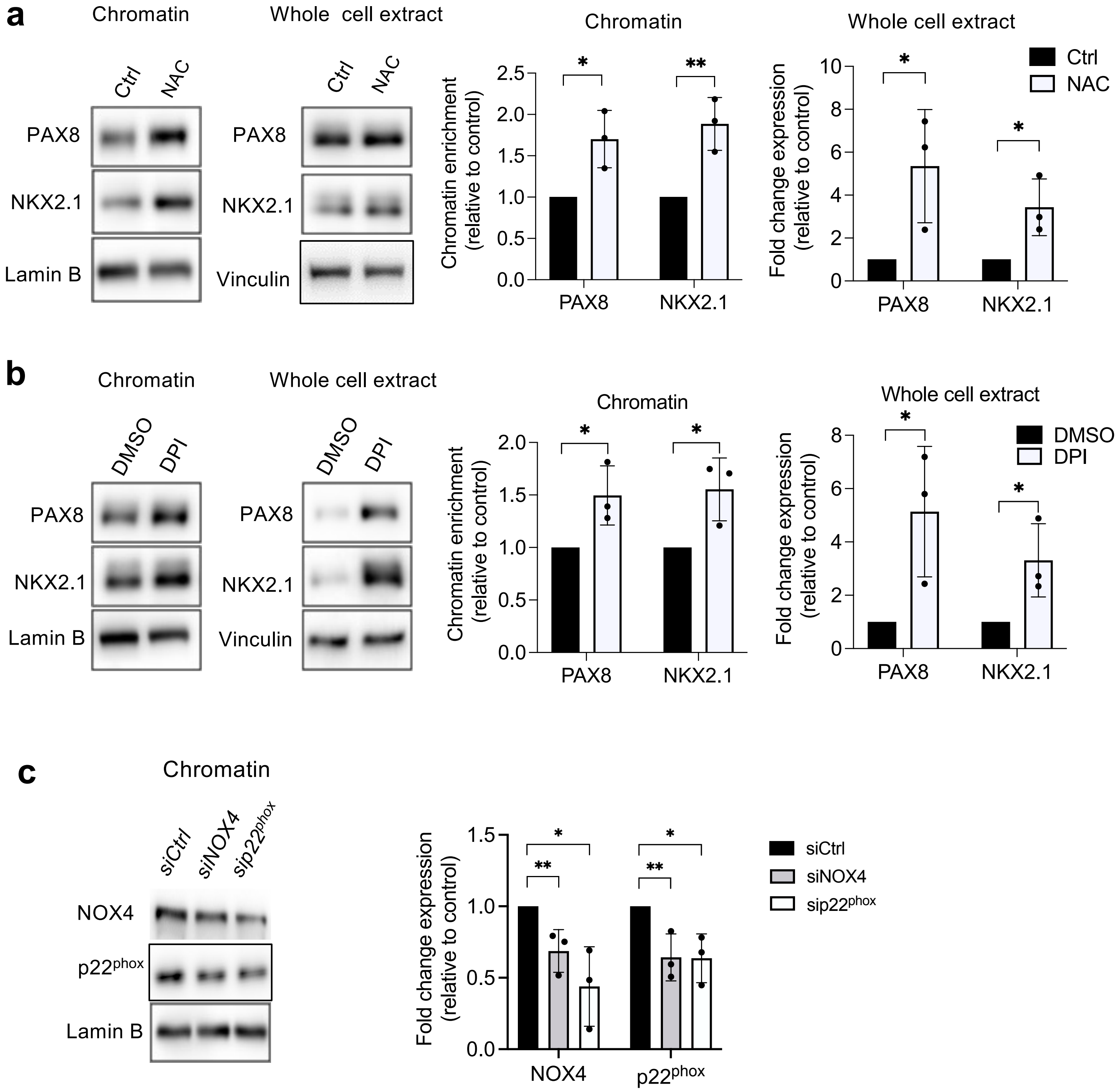


**Supplementary Figure 10**: Antioxidants promote recruitment of PAX8 and NKX2.1 to chromatin**. a**) BCPAP cells were pre-treated with 5 mM NAC for 2 h before being analysed for PAX8 and NKX2.1 protein expressions in chromatin fractions and whole-cell extracts. **b**) BCPAP cells were pre-treated with 1 μM DPI for 6 h before being analysed for PAX8 and NKX2.1 protein expressions in chromatin fractions and whole-cell extracts. c) siRNA knockdown validation of NOX4 or p22$^{phox}$ protein expression levels analysed by Western-blot in chromatin fraction 72h after knocking down of NOX4 or p22phox by RNA interference in BCPAP cells. Densitometry quantification of protein levels normalized to Lamin B or Vinculin levels and presented as chromatin enrichment or fold change compared with control cells. Values are mean ± SE. *p < 0.05 and **p < 0.01 (n = 3).

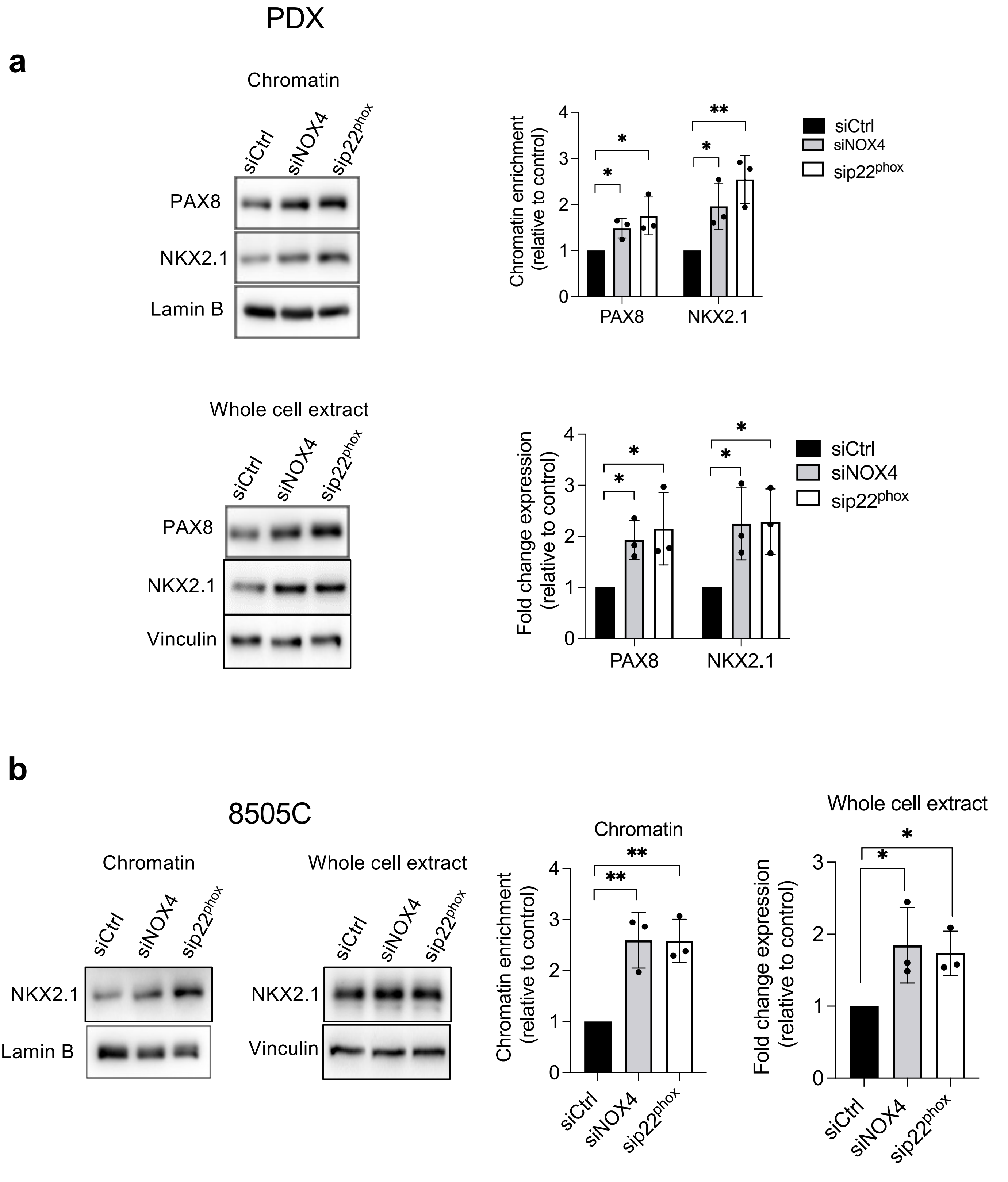


**Supplementary Figure 11**: NOX4 inhibits recruitment of PAX8 and NKX2.1 to chromatin. a) Western blot analysis of PAX8 and NKX2.1 protein expressions in chromatin fractions and whole-cell extracts 72 h after knocking down of NOX4 or p22phox by RNA interference in PDX cells. b) Western blot analysis of NKX2.1 protein expressions in chromatin fractions and whole-cell extracts 72 h after knocking down of NOX4 or p22phox by RNA interference in 8505C cells Densitometry quantification of protein levels normalized to Lamin B or Vinculin levels and presented as chromatin enrichment or fold change compared with control cells. Values are mean ± SE. *$p < 0.05$ and **$p < 0.01$ ($n = 3$).

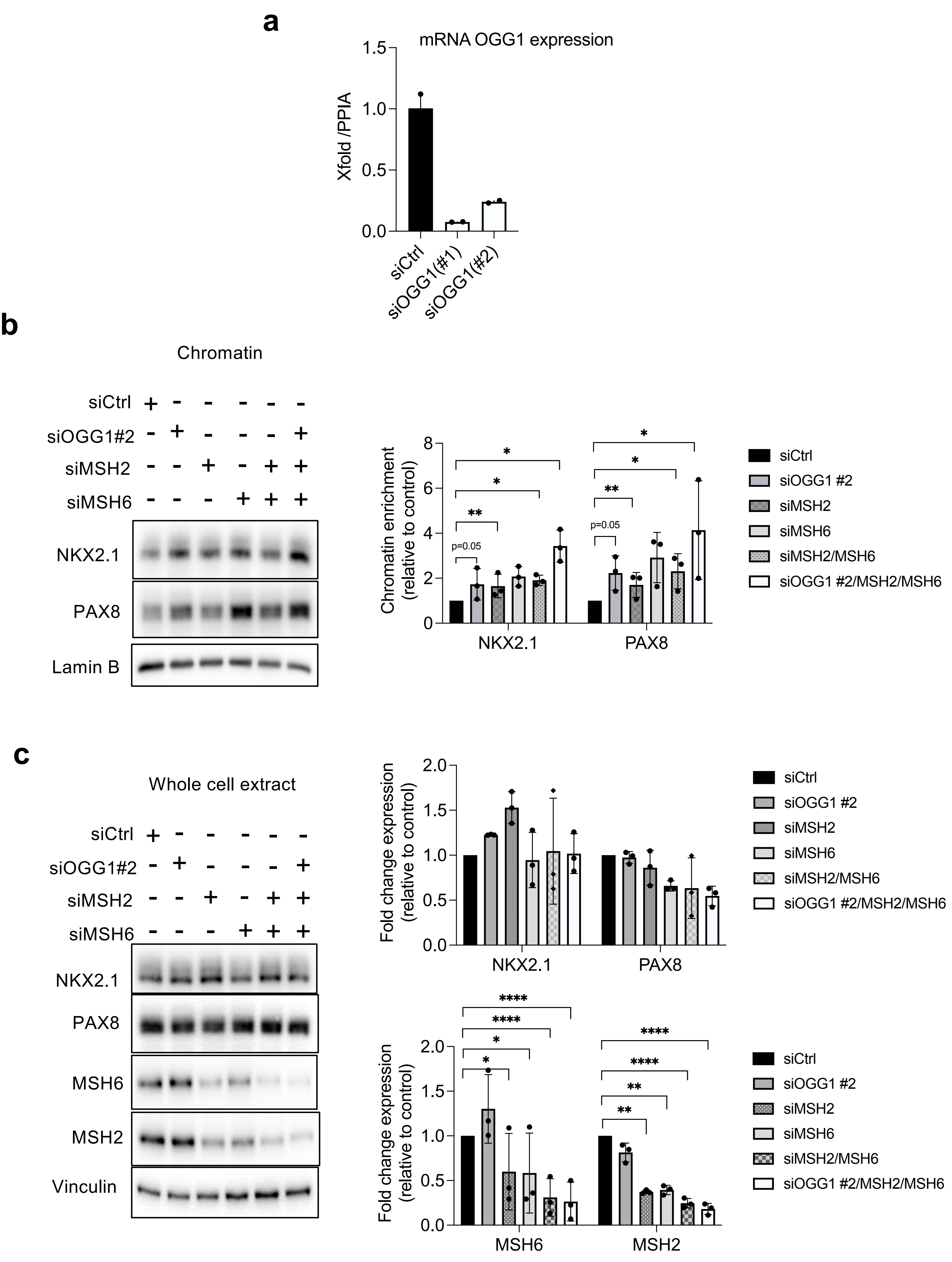
a
mRNA OGG1 expression
Xfold /PPIA
1.5
1.0
0.5
0.0
siCtrl
siOGG1(#1)
siOGG1(#2)
b
Chromatin
siCtrl
siOGG1#2
siMSH2
siMSH6
NKX2.1
PAX8
Lamin B
Chromatin enrichment (relative to control)
p=0.05
**
*
NKX2.1
PAX8
siCtrl
siOGG1 #2
siMSH2
siMSH6
siMSH2/MSH6
siOGG1 #2/MSH2/MSH6
c
Whole cell extract
siCtrl
siOGG1#2
siMSH2
siMSH6
NKX2.1
PAX8
MSH6
MSH2
Vinculin
Fold change expression (relative to control)
2.0
1.5
1.0
0.5
0.0
NKX2.1
PAX8
siCtrl
siOGG1 #2
siMSH2
siMSH6
siMSH2/MSH6
siOGG1 #2/MSH2/MSH6
****
*
**
MSH6
MSH2

**Supplementary Figure 12**: OGG1, MSH2, and MSH6 inhibit recruitment of PAX8 and NKX2.1 to chromatin in BCPAP cells. **a**) Validation of the siRNA transfection efficiency – Quantitative RT-PCR shows gene downregulation after 24h of transfection with two different siRNA targeting OGG1 (n=2) **b**) Western blot analysis of PAX8 and NKX2.1 protein expressions in chromatin fractions 72 h after knocking down of OGG1 or MSH2 or MSH6 by RNA interference in BCPAP cells. **c**) Western blot analysis of PAX8, NKX2.1, MSH6 and MSH2 protein expressions in whole-cell extracts 72 h after knocking down of OGG1 or MSH2 and MSH6 by RNA interference in BCPAP cells. Densitometry quantification of protein levels normalized to Lamin B or Vinculin levels and presented as chromatin enrichment or fold change compared with control cells. Values are mean ± SE. *$p < 0.05$ and **$p < 0.01$ and ****$p<0.0001$ ($n = 3$).

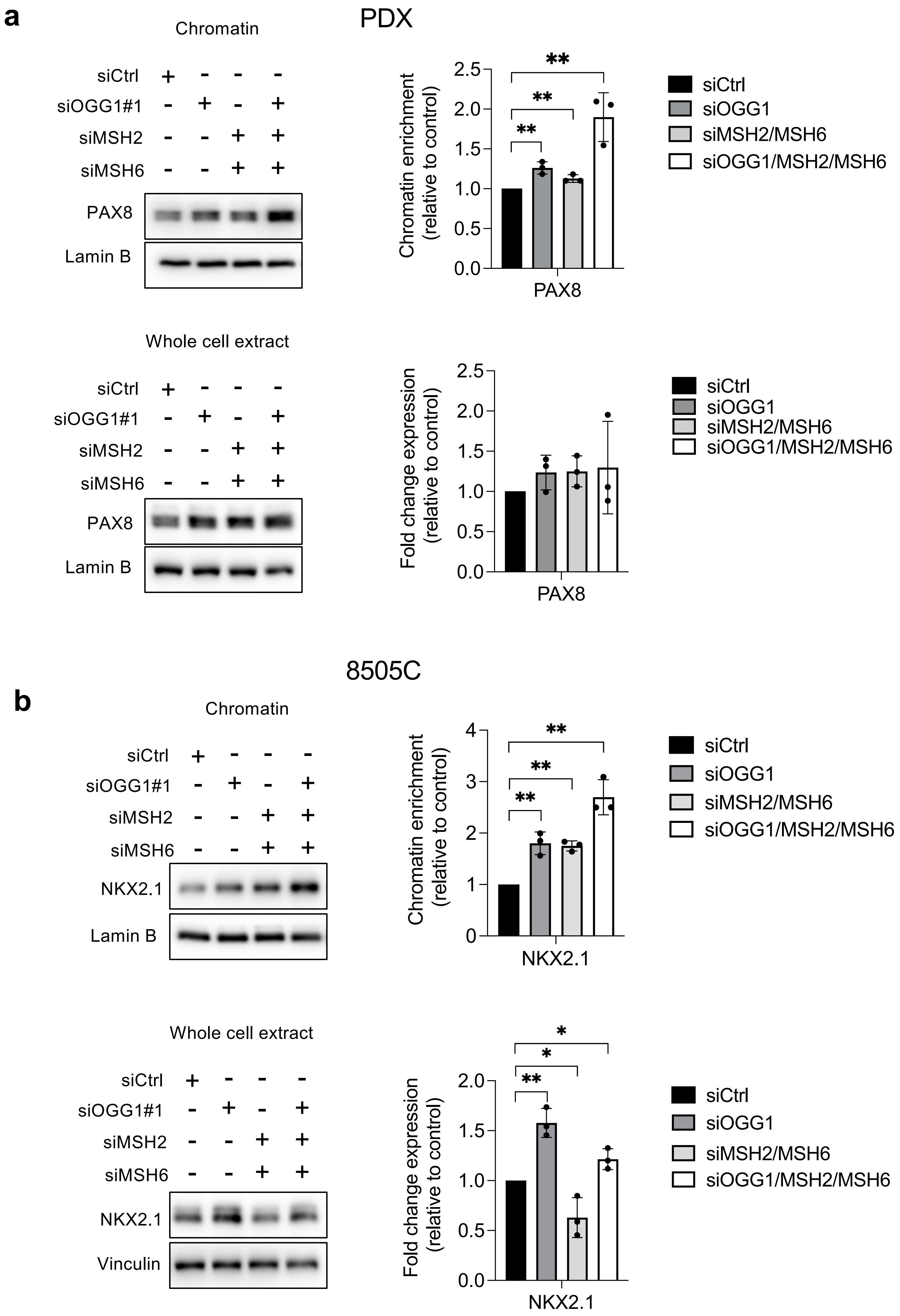
a
PDX
Chromatin
siCtrl
siOGG1#1
siMSH2
siMSH6
PAX8
Lamin B
Chromatin enrichment (relative to control)
2.5
2.0
1.5
1.0
0.5
0.0
**
**
**
PAX8
siCtrl
siOGG1
siMSH2/MSH6
siOGG1/MSH2/MSH6
Whole cell extract
siCtrl
siOGG1#1
siMSH2
siMSH6
PAX8
Lamin B
Fold change expression (relative to control)
2.5
2.0
1.5
1.0
0.5
0.0
PAX8
siCtrl
siOGG1
siMSH2/MSH6
siOGG1/MSH2/MSH6
8505C
b
Chromatin
siCtrl
siOGG1#1
siMSH2
siMSH6
NKX2.1
Lamin B
Chromatin enrichment (relative to control)
4
3
2
1
0
**
**
**
NKX2.1
siCtrl
siOGG1
siMSH2/MSH6
siOGG1/MSH2/MSH6
Whole cell extract
siCtrl
siOGG1#1
siMSH2
siMSH6
NKX2.1
Vinculin
Fold change expression (relative to control)
2.0
1.5
1.0
0.5
0.0
*
*
**
NKX2.1
siCtrl
siOGG1
siMSH2/MSH6
siOGG1/MSH2/MSH6

**Supplementary Figure 13**: OGG1, MSH2, and MSH6 inhibit recruitment of PAX8 and NKX2.1 to chromatin in PDX and 8505C cells. **a**) Western blot analysis of PAX8 protein expression in chromatin fractions and whole-cell extract 72 h after knocking down of OGG1 or MSH2 and MSH6 by RNA interference in PDX cells. **b**) Western blot analysis of NKX2.1 protein expression in chromatin fractions and whole-cell extracts 72 h after knocking down of OGG1 or MSH2 and MSH6 by RNA interference in 8505C cells. Densitometry quantification of protein levels normalized to Lamin B or Vinculin levels and presented as chromatin enrichment or fold change compared with control cells. Values are mean ± SE. *$p < 0.05$ and **$p < 0.01$ ($n = 3$).

**a**

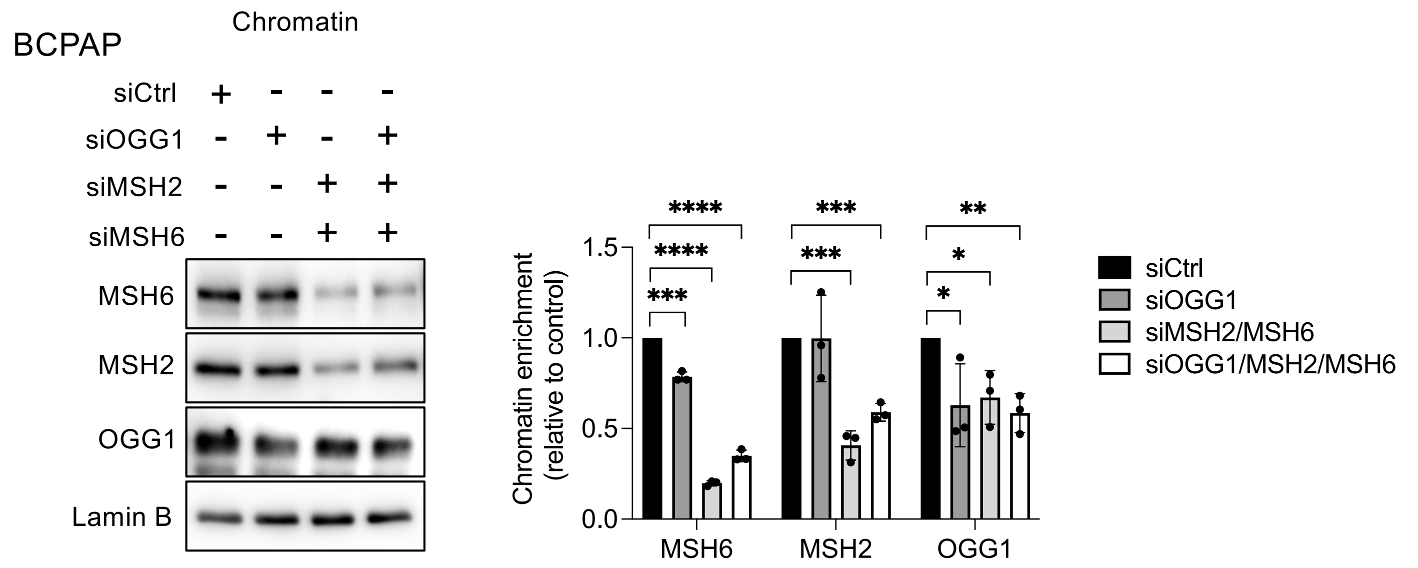

BCPAP
Chromatin
siCtrl + - - -
siOGG1 - + - +
siMSH2 - - + +
siMSH6 - - + +
MSH6
MSH2
OGG1
Lamin B
Chromatin enrichment (relative to control)
1.5
1.0
0.5
0.0
MSH6
MSH2
OGG1
****
****
***
***
***
**
*
*
siCtrl
siOGG1
siMSH2/MSH6
siOGG1/MSH2/MSH6


**b**

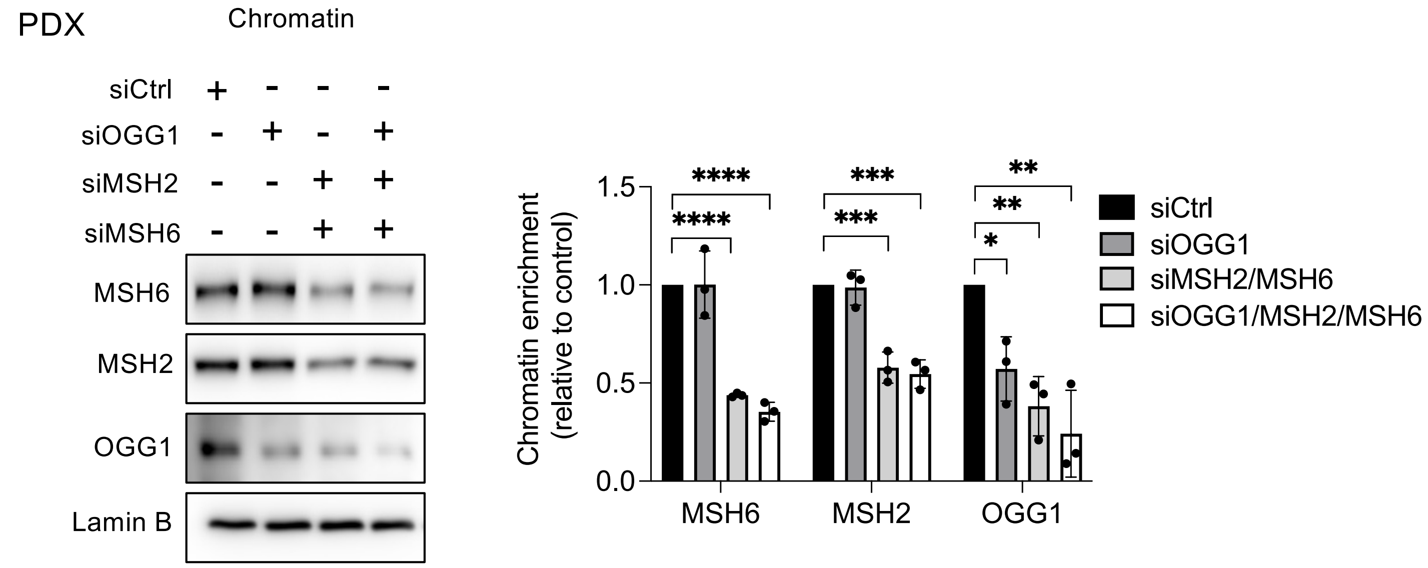

PDX
Chromatin
siCtrl + - - -
siOGG1 - + - +
siMSH2 - - + +
siMSH6 - - + +
MSH6
MSH2
OGG1
Lamin B
Chromatin enrichment (relative to control)
1.5
1.0
0.5
0.0
MSH6
MSH2
OGG1
****
****
***
***
**
**
*
siCtrl
siOGG1
siMSH2/MSH6
siOGG1/MSH2/MSH6


**c**

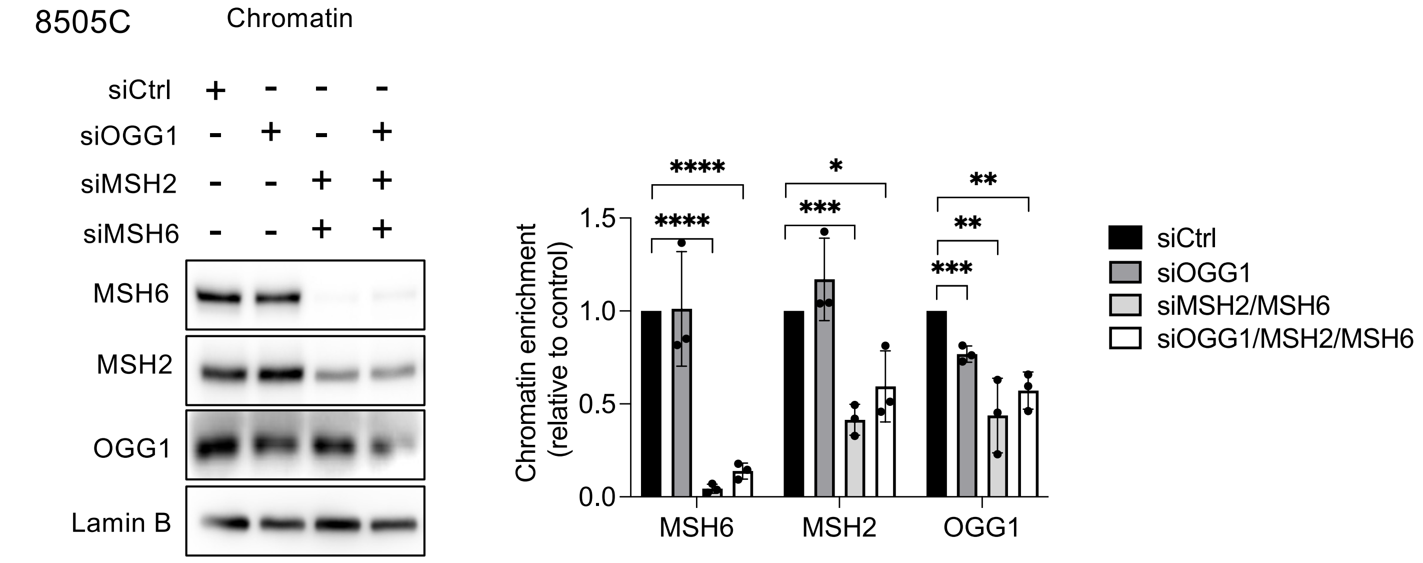

8505C
Chromatin
siCtrl + - - -
siOGG1 - + - +
siMSH2 - - + +
siMSH6 - - + +
MSH6
MSH2
OGG1
Lamin B
Chromatin enrichment (relative to control)
1.5
1.0
0.5
0.0
MSH6
MSH2
OGG1
****
****
*
***
**
**
***
siCtrl
siOGG1
siMSH2/MSH6
siOGG1/MSH2/MSH6

**Supplementary Figure 14**: siRNA knockdown validation of OGG1, MSH2, and MSH6 analyzed in chromatin fraction of BRAF-mutated thyroid cells. **a**) Western blot analysis of OGG1, MSH2, and MSH6 protein expressions in chromatin fractions 72 h after knocking down of OGG1, MSH2, and MSH6 in BCPAP cells. **b**) Western blot analysis of OGG1, MSH2, and MSH6 protein expressions in chromatin fractions 72 h after knocking down of OGG1, MSH2, and MSH6 in PDX cells. **c**) Western blot analysis of OGG1, MSH2, and MSH6 protein expressions in chromatin fractions 72 h after knocking down of OGG1, MSH2, and MSH6 in 8505C cells. Densitometry quantification of protein levels normalized to Lamin B levels and presented as chromatin enrichment compared with control cells. Values are mean ± SE. $^{*}p < 0.05$, $^{**}p < 0.01$, $^{***}p < 0.001$ and $^{****}p<0.0001$ ($n = 3$).

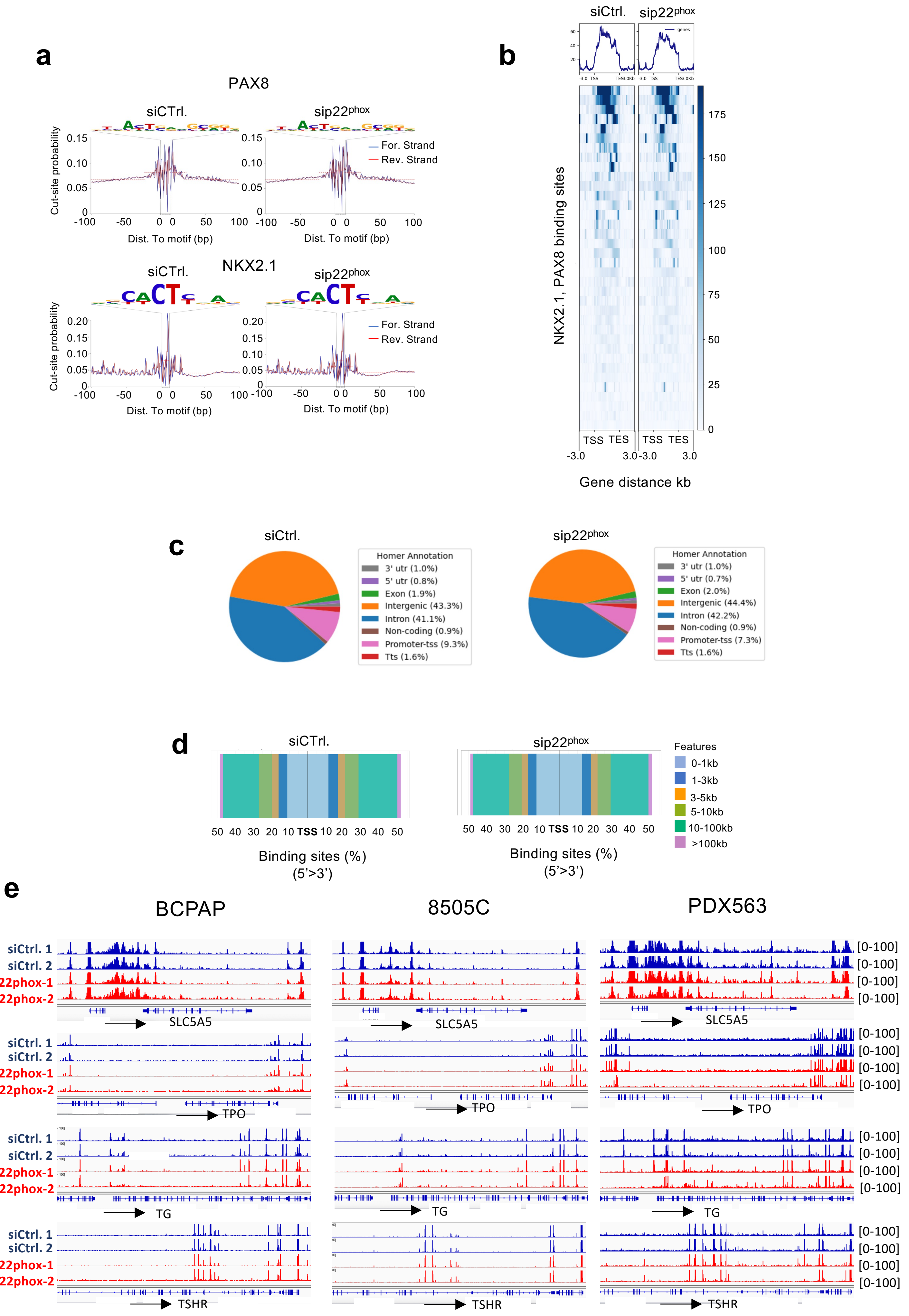
a
PAX8
siCTrl.
sip22phox
Cut-site probability
0.15
0.10
0.05
0
-100
-50
0
50
100
Dist. To motif (bp)
For. Strand
Rev. Strand
NKX2.1
0.20
b
siCtrl.
sip22phox
genes
NKX2.1, PAX8 binding sites
TSS
TES
-3.0
3.0
Gene distance kb
175
150
125
100
75
50
25
c
Homer Annotation
3' utr (1.0%)
5' utr (0.8%)
Exon (1.9%)
Intergenic (43.3%)
Intron (41.1%)
Non-coding (0.9%)
Promoter-tss (9.3%)
Tts (1.6%)
5' utr (0.7%)
Exon (2.0%)
Intergenic (44.4%)
Intron (42.2%)
Promoter-tss (7.3%)
d
Binding sites (%)
(5'>3')
Features
0-1kb
1-3kb
3-5kb
5-10kb
10-100kb
>100kb
e
BCPAP
8505C
PDX563
siCtrl. 1
siCtrl. 2
sip22phox-1
sip22phox-2
[0-100]
SLC5A5
TPO
TG
TSHR

**Supplementary Figure 15**: ATAC-seq analysis of BRAF-mutated thyroid cells deleted or not for $p22^{phox}$. **a**) ATAC-seq footprint at the PAX8 and NKX2.1 full sites. Cut sites probability, insertions per site are normalized to have the same average depth of insertions ± 100 bp away from motif center. The DNA sequences of the bottom are the motif of PAX8 and NKX2.1, respectively. **b**) Tornado plots of ATAC seq signals representing sites around the NKX2.1 and PAX8 genes (±3 kb center). **c**) Genomic distribution of ATAC-seq peaks in BCPAP cells transduced for 72 h with siRNA control and siRNA $p22^{phox}$. **d**) Distribution of transcription factor-binding loci relative to TSS. **e**) Representative sequencing tracks for the *SLC5A5, TPO, TG,* and *TSHR* loci show distinct ATAC-seq peaks. The ATAC-seq data have been normalized to take sequencing depth into account and the scale on the y-axis was chosen for optimal visualization of peaks for each sample (n = 2).

**8505C**

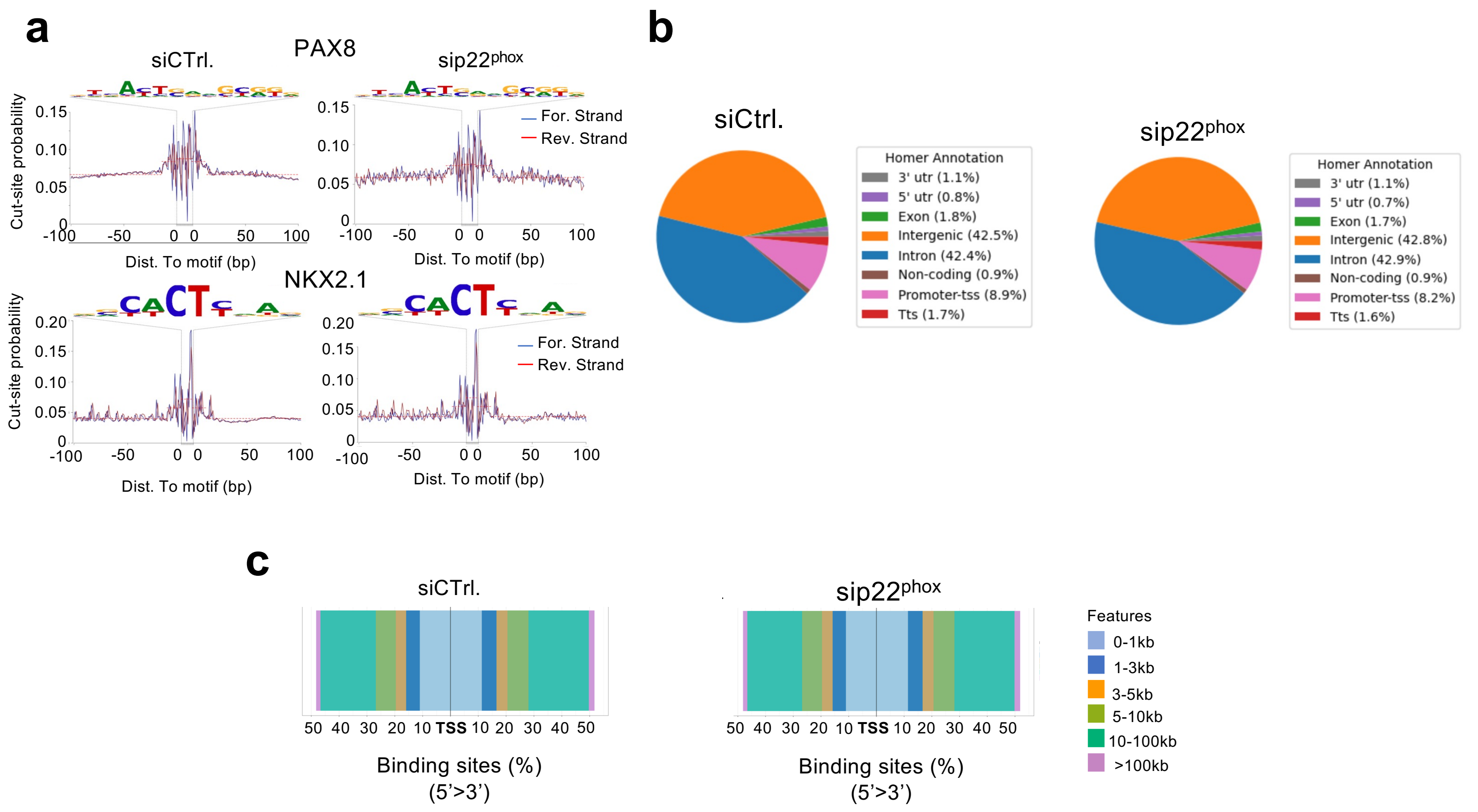
a
PAX8
siCTrl.
sip22phox
Cut-site probability
For. Strand
Rev. Strand
Dist. To motif (bp)
NKX2.1
b
siCtrl.
sip22phox
Homer Annotation
3' utr (1.1%)
5' utr (0.8%)
Exon (1.8%)
Intergenic (42.5%)
Intron (42.4%)
Non-coding (0.9%)
Promoter-tss (8.9%)
Tts (1.7%)
Homer Annotation
3' utr (1.1%)
5' utr (0.7%)
Exon (1.7%)
Intergenic (42.8%)
Intron (42.9%)
Non-coding (0.9%)
Promoter-tss (8.2%)
Tts (1.6%)
c
siCTrl.
sip22phox
50 40 30 20 10 TSS 10 20 30 40 50
Binding sites (%)
(5'>3')
Features
0-1kb
1-3kb
3-5kb
5-10kb
10-100kb
>100kb

**PDX**

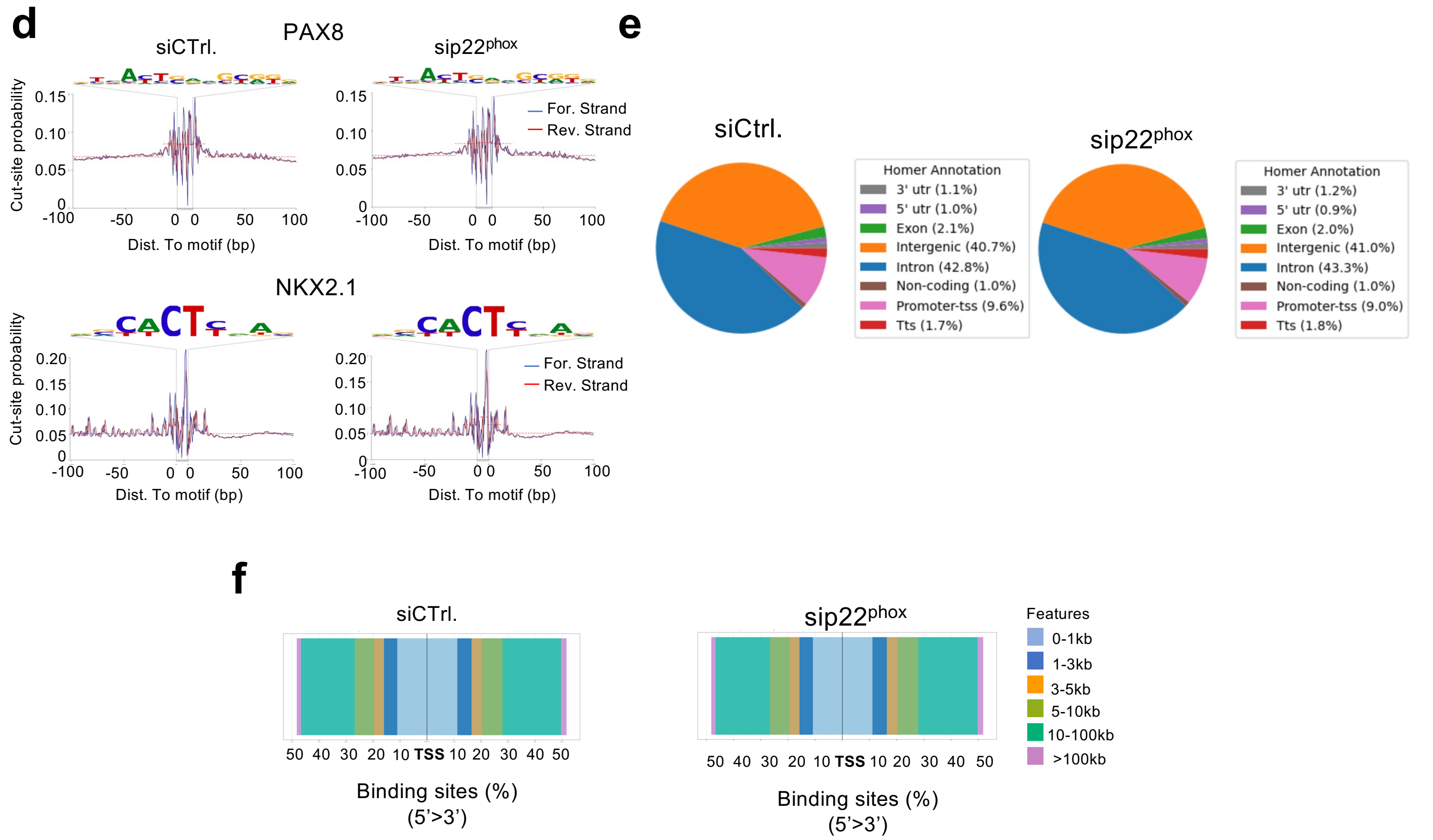
d
PAX8
siCTrl.
sip22phox
Cut-site probability
For. Strand
Rev. Strand
Dist. To motif (bp)
NKX2.1
e
siCtrl.
sip22phox
Homer Annotation
3' utr (1.1%)
5' utr (1.0%)
Exon (2.1%)
Intergenic (40.7%)
Intron (42.8%)
Non-coding (1.0%)
Promoter-tss (9.6%)
Tts (1.7%)
Homer Annotation
3' utr (1.2%)
5' utr (0.9%)
Exon (2.0%)
Intergenic (41.0%)
Intron (43.3%)
Non-coding (1.0%)
Promoter-tss (9.0%)
Tts (1.8%)
f
siCTrl.
sip22phox
50 40 30 20 10 TSS 10 20 30 40 50
Binding sites (%)
(5'>3')
Features
0-1kb
1-3kb
3-5kb
5-10kb
10-100kb
>100kb

**Supplementary Figure 16**: ATAC-seq analysis of 8505C and PDX cells deleted or not for p22$^{phox}$. **a, d**) ATAC-seq footprint at the PAX8 and NKX2.1 full sites. Cut sites probability, insertions per site are normalized to have the same average depth of insertions ± 100 bp away from motif center. The DNA sequences of the bottom are the motif of PAX8 and NKX2.1, respectively. **b, e**) Genomic distribution of ATAC-seq peaks in BCPAP cells transduced for 72 h with siRNA control and siRNA p22$^{phox}$. **c, f**) Distribution of transcription factor-binding loci relative to TSS (n = 2).

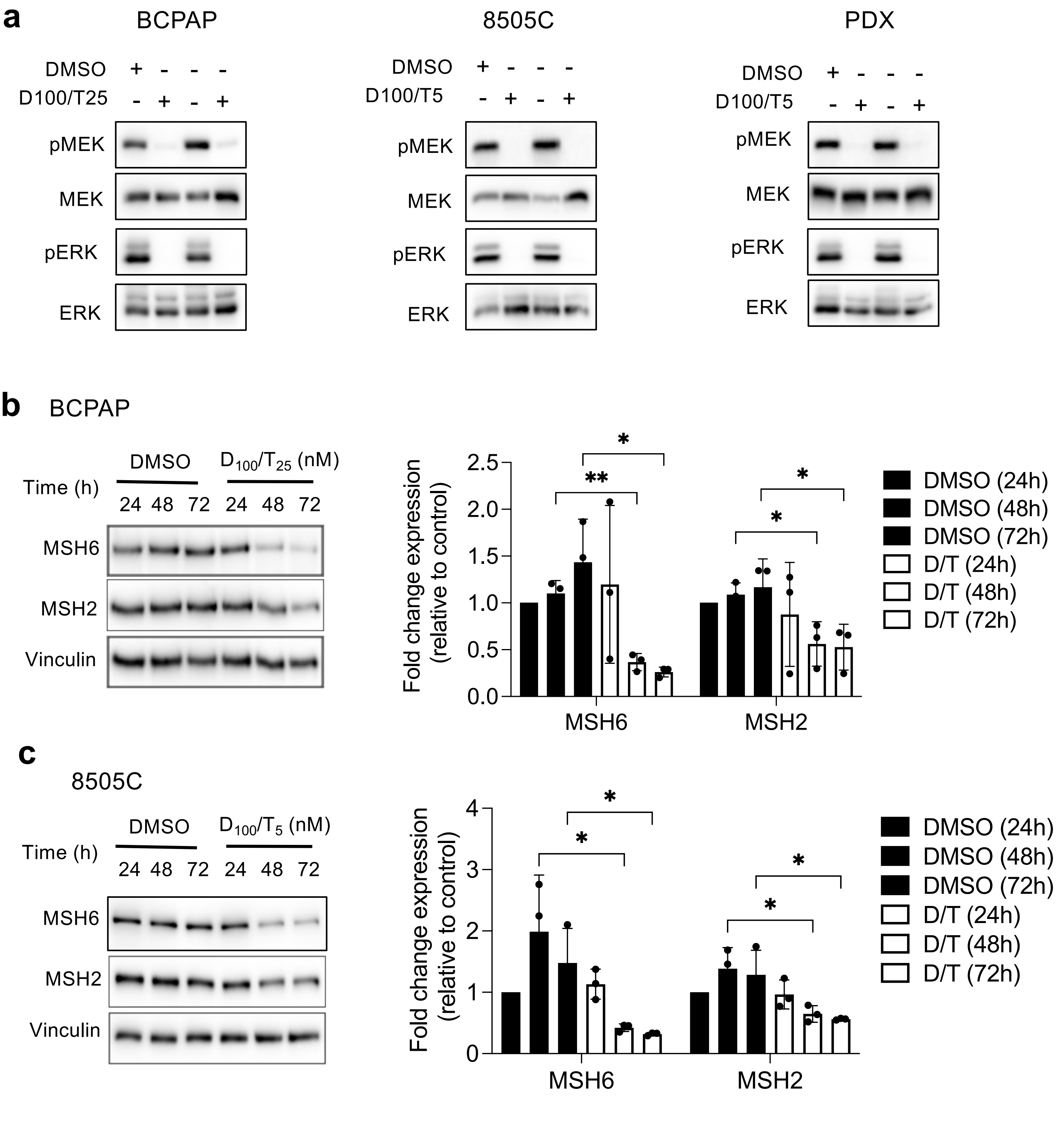

a
BCPAP
8505C
PDX
DMSO
D100/T25
D100/T5
pMEK
MEK
pERK
ERK
b
BCPAP
DMSO
D100/T25 (nM)
Time (h)
24 48 72 24 48 72
MSH6
MSH2
Vinculin
Fold change expression (relative to control)
MSH6
MSH2
DMSO (24h)
DMSO (48h)
DMSO (72h)
D/T (24h)
D/T (48h)
D/T (72h)
c
8505C
D100/T5 (nM)

**Supplementary Figure 17**: BRAF$^{V600E}$ regulates MSH2 and MSH6 expressions. **a**) BRAF-mutated thyroid cells were treated with Dabrafenib (100 nM) plus Trametinib (25 nM or 5 nM) for 4 h and analysed by Western-blot for expression of pMEK, MEK, pERK, ERK and pERK. **b**) Immunoblot detection of MSH2 and MSH6 in PDX cells after treatment by the combination of Dabrafenib plus Trametinib. **c**) Immunoblot detection of MSH2 and MSH6 in 8505C cells after treatment by the combination of Dabrafenib plus Trametinib. Densitometric quantification of protein levels normalized to vinculin levels and presented as fold change compared with vehicle-treated cells. The Student t-test was done by comparing combination versus DMSO for each corresponding time of kinetic. Values are mean ± SE. $*p < 0.05$ and $**p < 0.01$ ($n = 3$).

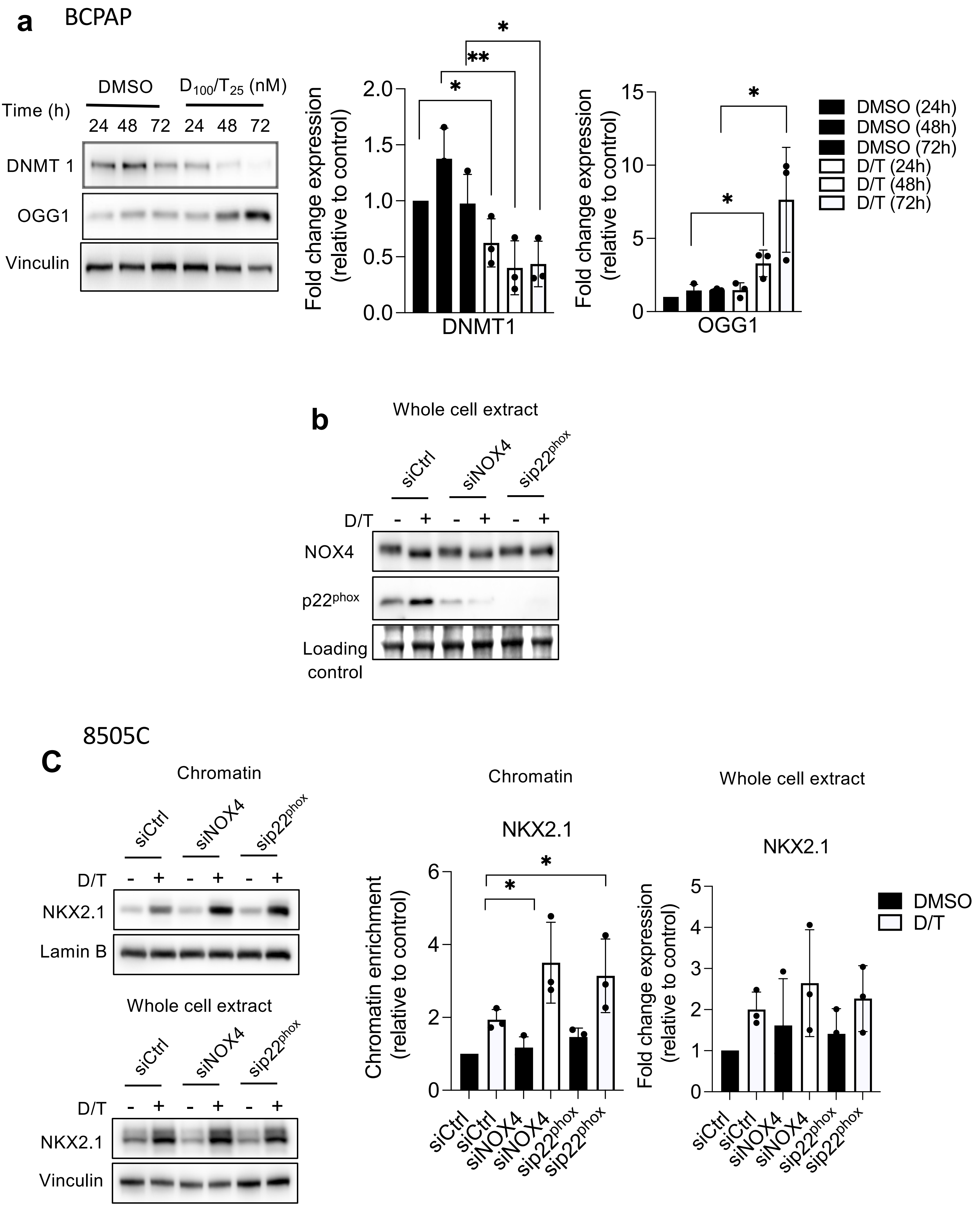

a
BCPAP
DMSO
$D_{100}/T_{25}$ (nM)
Time (h)
24 48 72 24 48 72
DNMT 1
OGG1
Vinculin
Fold change expression (relative to control)
DNMT1
OGG1
DMSO (24h)
DMSO (48h)
DMSO (72h)
D/T (24h)
D/T (48h)
D/T (72h)
b
Whole cell extract
siCtrl
siNOX4
$sip22^{phox}$
D/T
NOX4
$p22^{phox}$
Loading control
8505C
C
Chromatin
NKX2.1
Lamin B
Whole cell extract
Vinculin
Chromatin enrichment (relative to control)
DMSO
D/T

**Supplementary Figure 18**: Dabrafenib plus Trametinib combination increases OGG1 protein expression level in BRAF-mutated thyroid cells. **a**) Immunoblot detection of OGG1 and DNMT1 in BCPAP cells after treatment by the combination of Dabrafenib plus Trametinib. Densitometric quantification of protein levels normalized to vinculin levels and presented as fold change compared with vehicle-treated cells. The Student t-test was done by comparing combination versus DMSO for each corresponding time of kinetic. Values are mean ± SE. *$p < 0.05$, and **$p < 0.01$,. ($n = 3$). **b**) siRNA knockdown validation of NOX4 and $p22^{phox}$ protein expression levels analysed by Western-blot in whole-cell extract 72 h after knocking down of NOX4 or $p22^{phox}$ by RNA interference in BCPAP cells. c) Western-blot analysis of NKX2.1 protein expressions in chromatin fractions and whole-cell extracts of 8505C cells transduced with siRNA control or siRNA NOX4 or siRNA $p22^{phox}$ and treated with Dabrafenib plus Trametinib combination for 48 h. Densitometric quantification of protein levels normalized to loading control and presented as fold change compared with siRNA control-transduced cells. Values are mean ± SE. *$p < 0.05$ ($n = 3$).

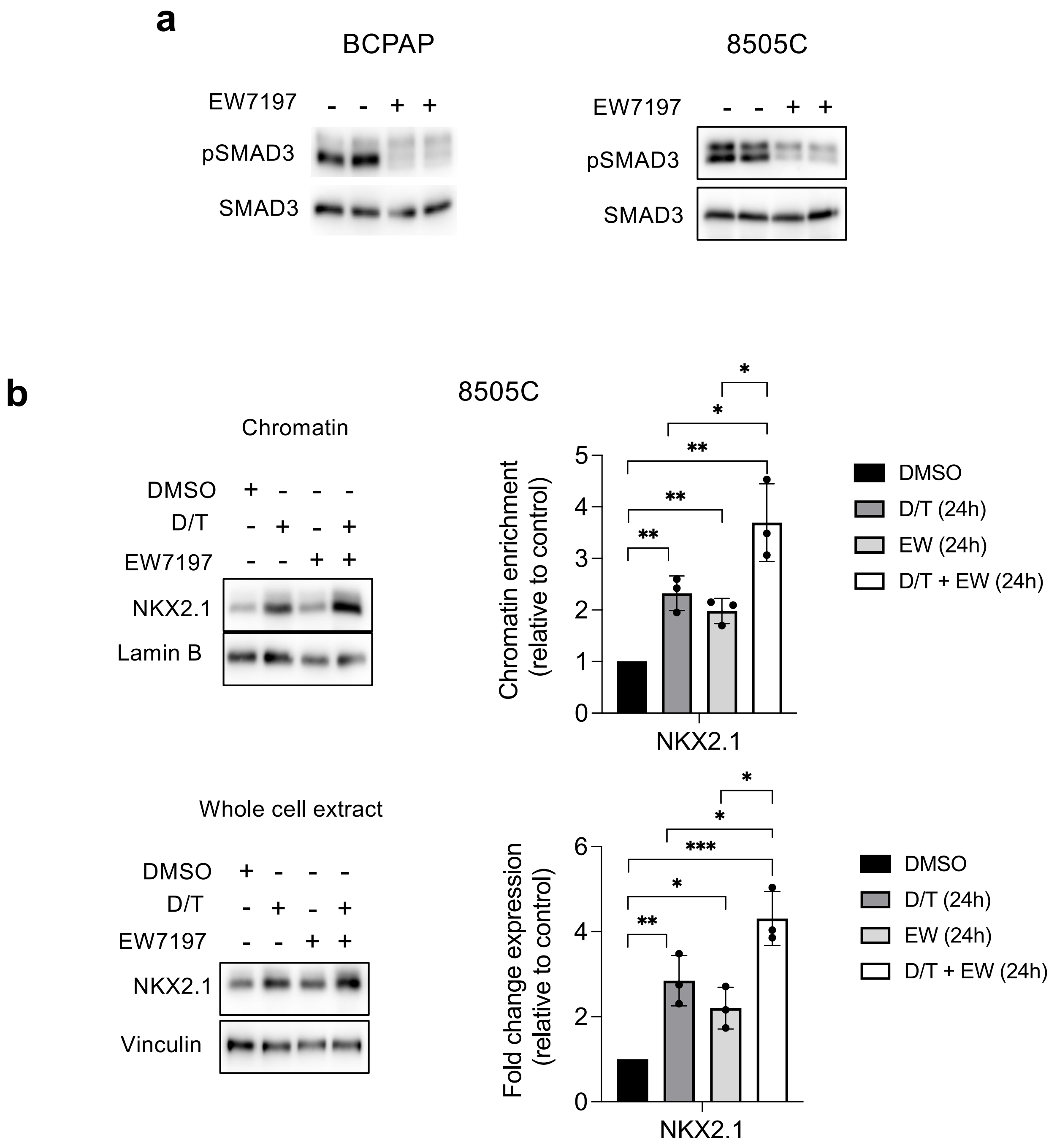


**Supplementary Figure 19**: Co-inhibition of SMAD and MAPK signaling promotes NKX2.1 recruitment to chromatin. **a**) BRAF-mutated thyroid cells were treated with TGF-beta receptor inhibitor (EW7197, 1 μM) for 4 h and analyzed by Western-blot for expression of pSMAD3 and SMAD3. **b**) Western-blot analysis of NKX2.1 protein expressions in chromatin fractions and whole-cell extracts of 8505C cells treated or not with Dabrafenib plus Trametinib combination in the presence or the absence of EW7197 for 24 h. Densitometry quantification of protein levels normalized to Lamin B or Vinculin levels and presented as chromatin enrichment or fold change compared with control cells. Values are mean ± SE. *$p < 0.05$, **$p < 0.01$, and ***$p < 0.001$ (n=3).
