## Supplementary Table for "NOX4 prevents the recruitment of PAX8 and NKX2.1 to chromatin in BRAF-mutated thyroid cancer cells"

**Supplementary Table 1** : Details of samples available for Immunohistochemistry analysis

| Sample ID | BRAF mut. | Tumor cell % | Tissu Type |
| --- | --- | --- | --- |
| 05-01 | p.V600E | 80 | Primitive Tumor |
| 04-16 | p.V600E | 90 | Lymph Node |
| 14-36 | p.V600E | 90 | Metastasis |
| 11-33 | p.V600E | 90 | Primitive Tumor |
| 02-02 | p.V600E | 90 | Primitive Tumor |
| 14-22 | p.V600E | 80 | Metastasis |
| 06-06 | p.V600E | 80 | Primitive Tumor |
| 01-23 | p.V600E | 90 | Metastasis |
| 14-37 | p.V600E | 90 | Primitive Tumor |
| 01-13 | p.V600E | 80 | Primitive Tumor |
| 06-28 | p.V600E | 90 | Primitive Tumor |
| 03-14 | p.V600E | 90 | Primitive Tumor |
| 01-35 | p.V600E | 90 | Primitive Tumor |
| 01-39 | p.V600E | 80 | Primitive Tumor |
| 03-15 | p.V600E | >90 | Metastasis |
